# Large-scale single-molecule analysis of tau proteoforms

**DOI:** 10.1101/2025.06.26.660445

**Authors:** J Joly, V Budamagunta, Z Zhang, B Nortman, M Jouzi, R Bhatnagar, JD Egertson, ME Flaster, R Grothe, S Guha, K Kaneshige, K McVey, N Nelson, RT Perera, SJ Tan, T Trinh, D Arnott, J Lipka, NJ Pandya, L Rougé, TJ Wendorff, DS Kirkpatrick, A Rohou, DC Butler, S Lotz, A Forton, ER Sartori, JE Schwarz, PS Brereton, K Chen, MA Darcy, HR Golnabi, R Hartley, PF Indermuhl, CE Inman, KM Jin, S Katsyuk, R Kota, B Lowry, JC Menger, DA Miller, MR Newman, A Ogunyemi, JK Robinson, N Steiner, J Sun, S Tabakman, L Wang, Z Wang, SK Wilcox, NAUTILUS BIOTECHNOLOGY, GT Kapp, S Patel, S Temple, T Bertucci, J Blanchard, A Huhmer, S Sankar, K Juneau, P Mallick

## Abstract

Proteins exist as diverse proteoforms resulting from a combination of genetic variation, alternative splicing, and post-translational modifications. Current methods struggle to capture this complexity at the single-molecule level. Here we introduce Iterative <u>Ma</u>pping of <u>P</u>roteoforms (IMaP), a method that enables the massively-parallel interrogation of millions to billions of single-protein molecules through iterative probing with fluorescently labeled antibodies. Using 12 site-specific antibodies, the method is capable of measuring 2^12^ (4,096) potential proteoform groups. We used IMaP to measure proteoform group profiles of the tau protein, a key player in neurodegenerative diseases, using two pan anti-tau antibodies (Tau-13, Tau-216), three isoform-specific antibodies (Anti-0N, Anti-2N, Anti-4R), and seven phosphosite-specific antibodies (Anti-pT181, Anti-pS202+pT205, Anti-pT205, Anti-pS214, Anti-pT217, Anti-pT231, and Anti-pS396). The method demonstrates high sensitivity (detecting proteoforms at 0.1% abundance), high reproducibility (median CV <5.5%), and broad dynamic range (>3 orders of magnitude), outperforming conventional techniques in resolving closely related proteoform groups. We demonstrated that the method can be used on relevant biological samples by examining various neuronal models (iNeuron cells, organoids, MiBrains, and mouse brains) and human samples. This examination revealed 130 distinct tau proteoform groups with as many as six phosphorylation events. The non-random distribution of these phosphorylation events suggests ordered and site-specific modification processes rather than random, stochastic accumulation. Certain combinations of phosphorylation events were more abundant than others; for example, pT217 preferentially co-occurred with pT181. In validating the applicability of the assay to human disease samples, we noted a specific pattern of multiple phosphorylation events in an advanced Alzheimer’s disease patient that suggests a sequential pathway of pathological tau modification. Iterative Mapping of Proteoforms provides insights into proteoform complexity at the single-molecule level, with significant implications for understanding protein regulation in neurodegenerative diseases and beyond.

## INTRODUCTION

Cellular regulation is driven by a complex interplay between protein synthesis, transport, modification, and degradation. The most prevalent proteomics methods to measure these processes, shotgun mass spectrometry (MS) ^1^ and affinity-based profiling ^2 3^, focus on estimating a sample’s protein composition by measuring the abundance of peptide fragments or the binding of specific affinity reagents.

While these methods can also quantify peptides or individual sites that contain sequence variants or post-translational modifications (PTMs), they are unable to capture one of the most critical aspects of protein-based regulation—combinations of protein alterations. Specifically, a combination of genetic polymorphisms, RNA splice variants, proteolytic processing, and a multiplicity of PTMs may collectively generate millions of different proteoforms ^4^. Protein alterations can influence a protein’s structure ^5^, localization ^6^, interactions ^7^, stability ^8^, and enzymatic function ^7,9^. However, outside of a few specific examples ^10^, such as histones ^11^, very little is known about how proteoforms or a diverse set of proteoforms drive cellular function. Fundamental questions remain, including: ‘which of the near-infinite number of possible proteoforms exist?,’ ‘is there an order and timing by which a given protein molecule acquires multiple modifications?,’ and ‘how do molecularly heterogeneous sets of proteoforms work in concert to drive cell behavior?’

Despite the many unknowns, recent studies have established connections between proteoform alterations and complex biological processes ^12^. Proteoforms have also been implicated in cancers and neurodegenerative disorders ^13^. As a result, scientists increasingly emphasize the need to study proteins at the proteoform level rather than relying solely on measurements of protein abundance ^12^.

The current gap in understanding about the functional impacts of proteoforms arises because their measurement is extremely challenging. Proteoforms are typically present at low abundance, making them difficult to detect directly within complex biological matrices and requiring prefractionation or enrichment^14^. This challenge is compounded by the limitations of conventional separation techniques, which often fail to distinguish structurally similar proteoforms ^15^. Even when separated, mass spectrometry—despite its power—struggles to resolve isotopomers, in which PTMs occur at different sites but result in indistinguishable masses or when structural isomers have identical fragmentation patterns ^16^. These analytical hurdles are further exacerbated by the requirement for large amounts of highly purified material, which is rarely accessible for low abundance proteoforms in biological samples. Identification is also constrained by the lack of a comprehensive reference database of human proteoforms, making confident assignment difficult even when high-quality data are available ^12^. Together, these challenges hinder systematic proteoform discovery and quantification, slowing efforts to decipher their biological significance.

To date, the most prevalent approach to measure proteoforms is based on top-down mass spectrometry (TDMS), which analyzes intact proteins without enzymatic digestion. TDMS can identify sequence variations and discover novel PTMs and associated proteoforms ^13^. TDMS is highly effective for measuring the relative abundance of proteoforms, proteoform families (sets of proteoforms originating from a single gene), or proteoform groups (sets of related proteoforms that share a defined set of observed alterations). TDMS typically achieves this by analyzing ensembles of millions of molecules, but this approach results in reduced sensitivity because of signal dilution across multiple charge states for each isotopologue and lack of chromatographic resolution ^17^. Recently, single-ion MS methods ^18^ that enable the detection of proteoform families have been introduced^19^. However, they are not typically able to easily resolve proteoforms that are not separable by mass or ion mobility. Despite its significant value in proteomics research, the difficulty of analyzing larger proteins creates a discovery bias towards smaller proteins and the complexity of TDMS workflows significantly constrains its practicality for widespread application.

An emerging class of single-molecule methods are anticipated to advance proteoform measurement. Single molecule counting methods have theoretical benefits for being able to measure rare proteoforms extremely sensitively and possibly with simpler experimental workflows than currently achieved with TDMS. In addition, it may be possible to examine modifications existing at multiple sites across a protein ^20^. A number of single-molecule peptide sequencing methods cannot measure proteoforms as they rely on protein digestion, similar to shotgun mass spectrometry ^21,22^. However, two classes of methods have emerged for single-molecule analysis of proteoforms and proteoform groups: protein fingerprinting and nanopore-based sequencing. Protein fingerprinting methods have been shown to use single-molecule fluorescence resonance energy transfer (FRET) ^23^ to distinguish isoforms of alpha-Synuclein. Though this analysis enabled isoform-specific identification and quantification in defined mixtures, it has not yet been applied for biological inquiry. In addition, it is unclear how it might examine a multiplicity of PTMs within the same protein molecule. Nanopore-based protein sequencing technologies ^24^ aim to identify and characterize full-length proteoforms. Recent nanopore technology improvements may allow for sequencing of closely-spaced phosphorylation in individual reads of single molecules and improved identification by rereading individual protein molecules, providing proof-of concept for a platform for highly accurate protein barcode sequencing and full-length proteoform identification ^25^. However, detection of complex and highly charged PTMs remains technically challenging and underexplored. While early nanopore protein sequencing prototypes exist, single-molecule library preparation methods from biological samples that are compatible with nanopore sequencing have not yet been demonstrated. It also remains unclear whether current nanopore systems provide the scale and throughput needed to measure the millions to billions of molecules required to capture the full dynamic range of biological systems.

Here we introduce and apply a new single-molecule method, which we call Iterative <u>Ma</u>pping of <u>P</u>roteoforms (IMaP), to overcome the key limitations of methods for quantifying proteoform groups. Iterative Mapping of Proteoforms is designed to reproducibly quantify pre-defined sets of proteoform groups across a wide range of abundances via massively parallel measurement of millions to billions of individual protein molecules.

We developed an assay, based upon the single-molecule Iterative Mapping of Proteoforms method, to measure tau proteoform groups encoded by the human *MAPT* gene. For simplicity, we refer to this “Iterative Mapping of Proteoforms of tau” assay below as the “tau proteoform assay.” Tau was chosen as a first exemplar given its critical importance in neurodegenerative diseases and its diverse landscape of splice variants and PTMs. Specifically, tau exists as six splicing isoforms in the adult human brain ^26,27^ and undergoes diverse PTMs including proteolysis ^28^, phosphorylation ^29^, acetylation ^30^, and ubiquitination ^29^, resulting in millions of potential tau proteoforms. Recently, the measurement of phosphorylated tau has been shown to have greater diagnostic power than measurements of total tau^31,32^. While much is known about tau, and its various splice-forms and PTMs, very little is known about its proteoform landscape, and how this varies in diverse biological systems.

We first evaluated the tau proteoform assay through a series of control samples of known composition and enriched biological lysates known to contain tau proteoforms associated with epitope-specific probes within in our assay. Next, we demonstrated that the assay can be applied to biological samples by measuring the complex landscape of tau proteoform groups in neuronal tissue model systems of tauopathies, including fronto-temporal dementia (FTD) and Alzheimer’s disease (AD). We resolved isoform-specific variation in maturing tau across model systems. We additionally identified which combinations of phosphorylation events are more frequently observed. To demonstrate that the assay can be applied to human-derived brain tissue samples, we analyzed a small number of samples from patients that were either cognitively normal or had AD-associated impairment. As part of our analysis, we examined if phosphorylation patterns are random or instead suggestive of a specific order of occurrence, finding evidence for both specificity and ordering. These measurements also potentially shed light on routes by which specific proteoforms are generated and provide a foundation for future research investigating which tau proteoforms contribute the most to neural dysfunction and neurodegeneration, and how.

## RESULTS

### Single-molecule Iterative Mapping of Proteoforms

The method we introduce here, Iterative <u>Ma</u>pping of <u>P</u>roteoforms (IMaP) (**Figure 1**), consists of four steps that are outlined in **Figure 1**. In the first step (**Figure 1A**), protein is extracted from tissue lysate, denatured, and functionalized with an amine-reactive methyltetrazine-PEG4-STP (4-Sulfo-2,3,5,6-tetrafluorophenyl) ester (mTz) click reagent. If needed, a sample can be enriched using protein-specific antibodies. Next, functionalized protein is conjugated to specialized nanoparticles that were designed and built using DNA-origami approaches^33^. At most one protein molecule can be conjugated to each nanoparticle. At no point is the protein digested. Next (**Figure 1B**), protein-nanoparticle conjugates are deposited into a lane of a custom microfluidics flow cell with a functionalized glass surface containing millions to billions of optically-resolvable landing pads, as previously described^34^. As the DNA-origami particles are fluorescently labeled, we can determine which coordinates of the array contain a protein-nanoparticle conjugate and which are empty by imaging in the 488-wavelength. In the final step, single molecules are iteratively probed in a massively parallel manner using a series of site-specific fluorescently labeled ^35^ antibodies (**Figure 1C**). Each iteration consists of a bind, rinse, image, and remove cycle. Specifically, a fluorescently labeled antibody probe is introduced into the flow cell and allowed to incubate. Next, unbound antibody is rinsed from the flow cell and the flow cell is optically imaged ^34^ to determine where the antibody has bound. Following imaging, bound antibodies are removed from the flow cell, preparing the flow cell for a subsequent iteration. As fluorescent images are acquired, positions of each single-molecule binding event are recorded. Collectively, probing of the single-molecule array produces a series of binary bind/no-bind outcome patterns (single molecule binding traces), which are used to determine the potential identity of the proteoform at each coordinate in the flow cell. The cumulative relative binding across a typical experiment is shown across 36 cycles in Supp. Fig 1. Note that the binding signal is stable across the 36 cycles and that there is no significant build-up of background signal. By counting the observed binding patterns and correcting for antibody false positive and false negative binding, we estimate the quantity of each proteoform within each sample (**Figure 1D**). Proteoform abundances are then reported in parts per million. We note that Iterative Mapping is a fully general method that may have diverse applications beyond the proteoform analysis described herein.

**Figure 1:**
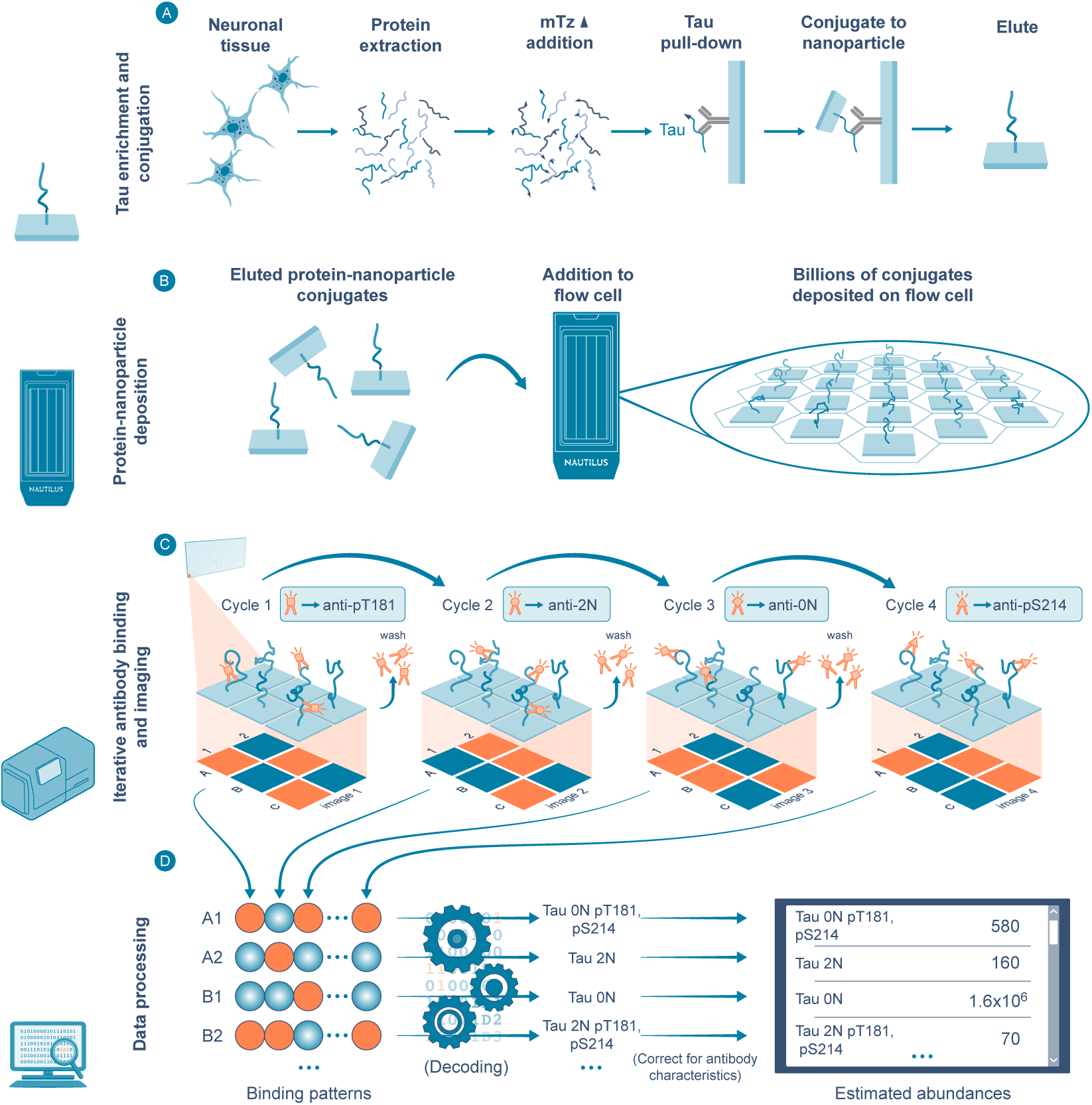
Single-molecule Iterative Mapping of Proteoforms workflow **Figure 1A:** Proteins are functionalized on lysine residues with methyltetrazine-PEG4-STP (4-Sulfo-2,3,5,6-tetrafluorophenyl) ester, conjugated by a cycloaddition reaction to a DNA origami nanoparticle containing a single trans-cyclooctene (TCO) moiety. **Figure 1B:** Tau protein nanoparticle-conjugates are deposited as single proteins on a glass surface chip. Between 4 and 12 sample lanes are configured in a flow cell for an experiment. **Figure 1C:** A series of fluorescently labeled antibodies are used to iteratively probe each molecule via a series of incubation, rinse, imaging, and washing steps. **Figure 1D:** Proteoforms are determined from the pattern of antibody binding and quantified by counting. A normalizing computation (not shown) is used to account for false-positive and false-negative binding of antibodies.

### Iterative Mapping of Proteoforms resolves tau proteoforms

We applied the Iterative Mapping of Proteoforms method to build a tau proteoform assay incorporating 12 antibodies that included two pan anti-tau antibodies (Tau-13, Tau-216), three isoform-specific antibodies (Anti-0N, Anti-2N, Anti-4R), and seven phosphosite-specific antibodies (Anti-pT181, Anti-pS202+pT205, Anti-pT205, Anti-pS214, Anti-pT217, Anti-pT231, and Anti-pS396) spanning key isoforms and phosphorylations in tau (**Figure 2A, Supp. Table S3**). 1N isoforms were inferred by lack of binding of the 0N and 2N antibodies. 3R isoforms were inferred by lack of binding of the 4R antibody. Phosphorylation at pS202 was inferred by binding of the Anti-pS202+pT205 antibody and lack of binding of the Anti-pT205 antibody. All other forms and modifications were measured directly from observed binding. An assay with 12 antibodies potentially measures up to 2^12^ (4,096) proteoform groups. Because 0N and 2N cannot both bind concurrently, and because we restrict our focus here to full-length tau, a tau proteoform assay restricted to molecules that have not been truncated enables us to measure 768 (6*2^7^) potential proteoform groups (**Figure 2A, Supp. Table S3**) of full-length tau. Given the targeted nature of the assay, there may exist additional protein alterations beyond those measured. As such, the assay measures proteoform groups rather than full proteoforms. The antibodies used herein were extensively validated for on-target, off-target, and non-specific binding as described in **Methods**. We included four control samples within each experiment comprising phosphorylated peptides, non-phospho peptides, and full-length tau proteoforms to measure the on and off target binding of each probe within that experiment (**Methods**). Representative images and a set of derived binary single-molecule bound/not-bound traces from the assay are shown in **Figure 2B**. Estimates of proteoform group are extracted from these traces for each molecule. For example, ‘Proteoform 2’ in **Figure 2B** shows binding with antibodies targeting 0N, pT181, and pS396, thereby identifying the molecule at that position as part of the proteoform group defined by the 0N-3R isoform with a double phosphorylation at sites pT181 and pS396. As noted above, quantification is derived from the assay by counting up the identifications and normalizing for false-positive and false-negative binding as described in **Methods**.

**Figure 2:**
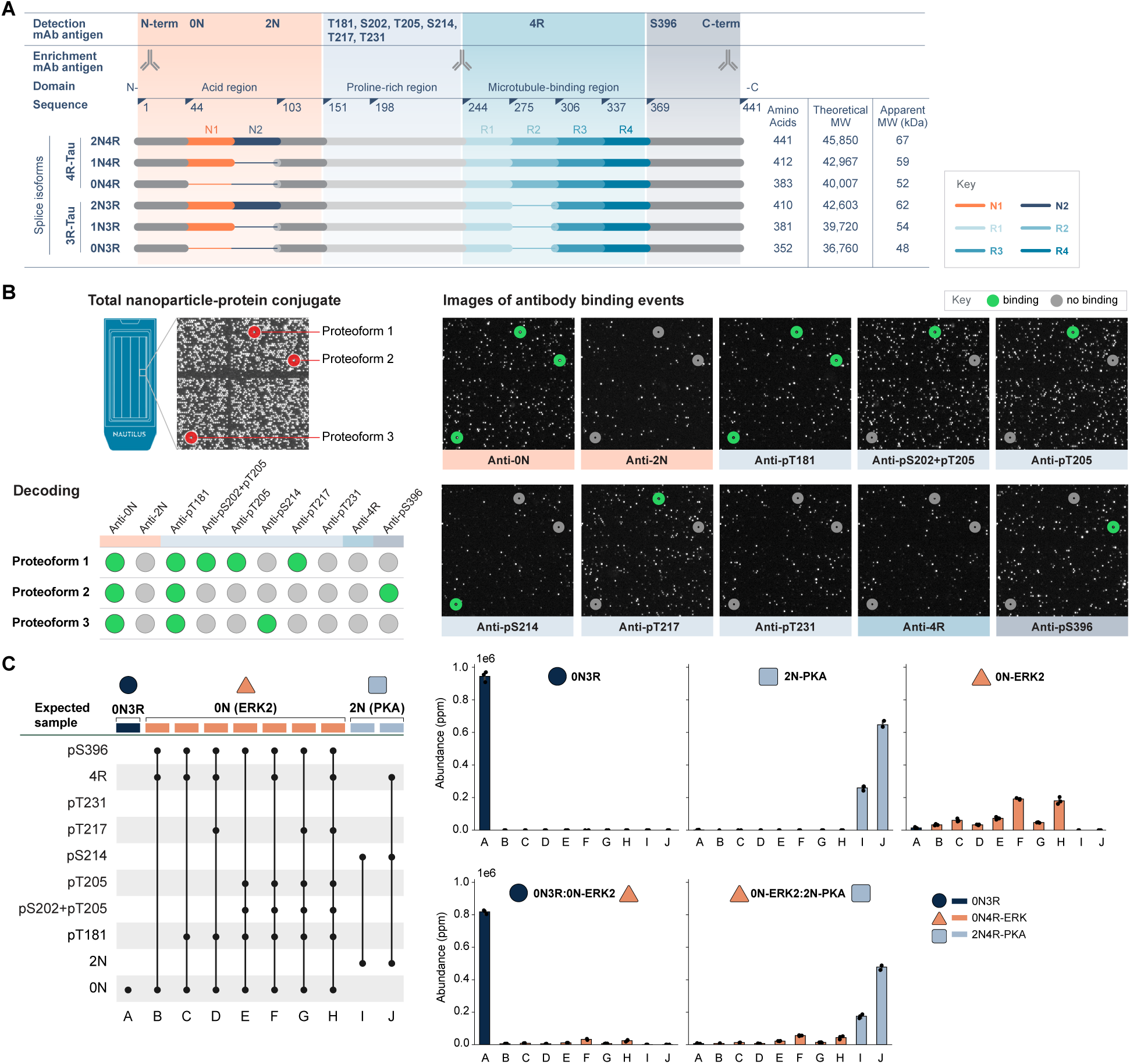
Application of iterative mapping to measure proteoforms of tau **Figure 2A:** Tau isoforms generated through tau gene expression, alternative RNA splicing, and protein translation. N1 and N2 tau isoforms (orange, dark navy blue) are generated by alternate splicing of exon 2 and 3 and alternate splicing of exon 10 produces the 3R and 4R isoforms (light blue). Approximate epitope positions for antibodies used for enrichment or for isoform and proteoform specific detection are shown. **Figure 2B:** Primary imaging data capturing a representative area of 130 µm x 130 µm of flow cell surface showing ∼16,000 landing pads. Each illuminated pixel is a single tau-nanoparticle-conjugate deposited in the hyper-dense array. Over a series of binding and imaging cycles a series of binding events from fluorescently labeled antibodies to individual protein molecules are recorded as traces giving information about the tau proteoform present at each coordinate. Green halo indicates positive binding event. **Figure 2C:** Tau proteoforms standards (0N3R, 2N4R-PKA, 0N4R-ERK) and expected epitope combination (A-J). The right panel shows the relative abundance of individual proteoforms in parts per million. The left panel shows the epitope profile of mixtures of tau proteoforms resolved by interactive mapping of proteoforms. The mixtures of protein standards are: 33.3% 0N-ERK2: 66.6% 0N3R and 33.3% 0N-ERK2: 66.6% 2N4R-PKA. NOTE: 2N4R-PKA represents tau proteoform 2N4R treated with Protein Kinase A, which is phosphorylated to >99.9% at Serine 214 (pS214), 0N-ERK2 denotes 0N4R treated with ERK2, giving rise to multiple phosphorylations. Error bars represent standard deviation and each sample has three technical replicates.

We developed a set of recombinant tau proteoform standards to enable characterization of assay performance. We specifically used recombinantly expressed and purified 0N4R and 2N4R isoforms of tau, and derivatives thereof (2N-PKA and 0N-ERK2) that had been phosphorylated by either protein kinase A (PKA) or Mitogen-activated protein kinase 1 (ERK2) (**Supp. Methods**). PKA preferentially modifies 2N4R at pS214 with approximately 99% completeness. ERK2 is less specific, dominantly modifying 0N4R at pT181, pS202/T205, and pS396 with approximately 90% phosphorylation efficiency (**Supp. Table S2**). Consequently, samples made with these standards contain a mixture of three dominant proteoforms and a series of proteoforms that arise from incomplete modification (**Figure 2C-left**). To mimic cell-derived tau, these standards can be mixed into a background cell-line that does not contain tau. Studies herein used Expi293F cells for spike-in experiments. The individual estimates of the composition of each standard are shown in the right panel in **Figure 2C**. Each standard has been assigned a color of either navy blue, orange, or light blue. These standards were mixed in a variety of ratios and assayed, demonstrating that the various proteoform groups were readily resolved (**Figure 2C-right)**.

To examine how this proteoform reference standard mixture can be measured by other common analytical approaches, we performed western blots (WB). As WB relies on electrophoretic separation and antibody visualization, we expected it to be less precise in resolving complex tau proteoforms. Analysis of an aliquot of a 1:2:3 mixture of three tau proteoforms—0N4R, 2N4R, and 2N4R-PKA—using silver staining successfully resolved the 0N and 2N isoforms (Supp. Fig. 2A**, lanes 2-3**). Detection with the pan-tau and isoform-specific antibodies confirmed the presence of the 0N4R and 2N4R tau isoforms (Supp. Fig. 2B) but was unable to differentiate the 2N4R from the 2N4R-PKA proteoform in the lane with a pure standard or in the mixtures. Similarly, the epitope-specific probe anti-pS214 specifically stained the 2N4R-PKA standard (but not the highly phosphorylated control standard 0N4R-ERK2 lacking pS214) as well as the protein standard mixtures (Supp. Fig. 2C). However, the specific proteoforms in the mixture and their relative phosphorylation site occupancy remained unresolved. These results affirm that WB provides limited resolution of tau isoforms, while the tau proteoform assay enables detailed proteoform characterization, including the detection of phosphorylated proteoforms in mixtures, establishing it as a higher resolution method for analyzing tau proteoforms.

### Accuracy, reproducibility, and dynamic range of the tau proteoform assay

We characterized the quantitative accuracy and reproducibility of the tau proteoform assay using mixtures of three tau protein standards (0N4R, 2N4R, and 2N4R-PKA) at molar ratios of 1:2:3 and 1:1:1. The mean absolute percent error for the 4R tau proteoforms was 9.97% (**Figure 3A**) across 21 replicates. We further examined reproducibility by analyzing the protein standard mixtures in a range of reproducibility challenges using a nested tree design. Overall, the median CV amongst technical replicates of prepared nanoparticle protein conjugates within the same experimental run was 1.5%. In addition to examining reproducibility within runs of the prepared control mixture within the same experiment, we examined the reproducibility across five batches of flow cells (CV=2.6%) and across three instruments (CV=3.6%). We additionally examined reproducibility of full end-to-end experiments that included library preparation performed 3X by the same operator (CV=5.5%) or as performed by three different operators (CV=5.5%). Though 90% of the intra-run variation ranges from 0-2.3%, percent, 90% of the interoperator variation ranges from 0 to 15.6%. Overall, the assay demonstrated strong reproducibility (**Figure 3B**) across a reproducibility challenge consisting of 297 sets of measurements.

**Figure 3:**
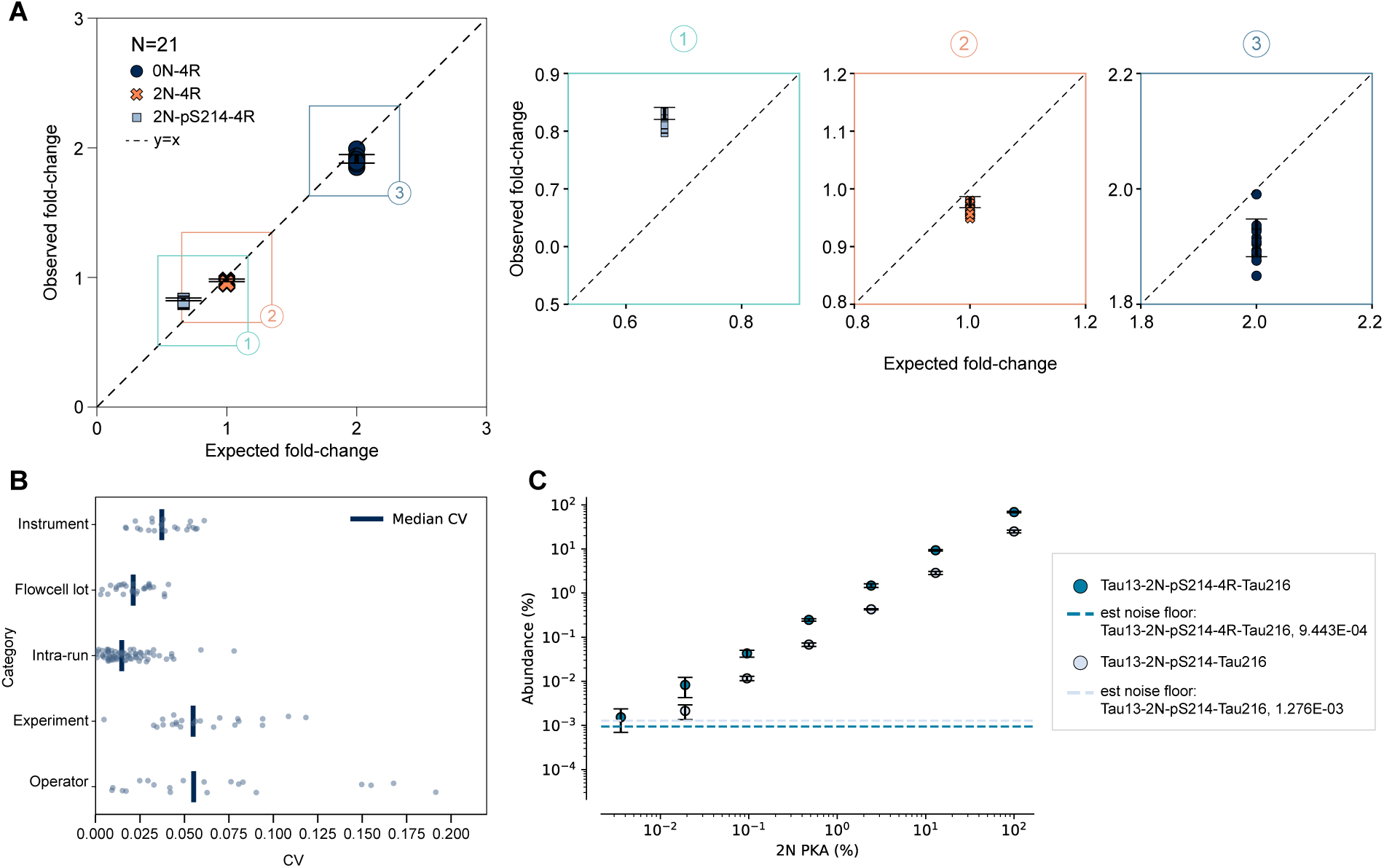
Accuracy, reproducibility, and dynamic range of the tau proteoform assay **Figure 3A.** Quantification accuracy and relative abundance were assessed using two mixtures of tau proteoform standards, each containing 0N4R, 2N4R, and 2N4R-PKA at defined molar ratios of 1:2:3 and 1:1:1, respectively. The results shown represent 21 technical replicates comparing the measured ratios between the two mixtures, with error bars indicating the standard deviation. The mean absolute percent error across all measurements was 9.97%. **Figure 3B.** Here we examined the reproducibility of quantitation. Overall, the median CV amongst technical replicates of prepared nanoparticle protein conjugates within the same experimental run was 1.5%. In addition to examining reproducibility within runs of the control mixture within the same experiment, we examined the reproducibility across batches of five flow cell batches (CV=2.6%) and across three instruments (CV=3.6%). We additionally examined reproducibility of full end-to-end experiments including library preparation performed by the same operator 3X (CV=5.5%) and as performed by three different operators (CV=5.5%). In total, estimates of reproducibility are drawn from 297 measurements. **Figure 3C:** Dynamic range was assessed using the 2N4R-PKA proteoform standard representing the proteoform 2N4R-pS214 with serial dilution into an orthogonal background of the 0N3R proteoform employing five (anti-Tau13, anti-2N, anti-pS214, anti-4R, anti-Tau216) detection antibodies (dark blue circle) and four (anti-Tau13, anti-2N, anti-pS214, anti-Tau-216) detection antibodies (light blue). The limit of quantitation (LOQ) was defined as the lowest point on the titration curve at which the signal to noise ratio is greater than 10 and the coefficient of variation is under 30%. Error bars represent the mean of nine technical replicates across three experiments.

Tau protein is relatively low in abundance in neuronal tissue, constituting approximately 0.09% to 0.7% of total protein ^36^. To address this, we developed a bead-based enrichment method using a combination of three distinct anti-tau antibodies (**Figure 1A, Suppl. Table S3**), which is compatible with the single-molecule library preparation workflow. Comparing proteoform abundances between the unenriched mixture and the enriched samples revealed a strong correlation (R² = 0.95, stddev 0.2%-1.03%), indicating consistent relative recovery across proteoforms with minimal bias against the specific tau species tested (**Suppl. Fig. 3**).

We established the dynamic range and linearity of tau proteoform quantification using spike-in experiments. In this context, dynamic range refers to the quantification of decreasing percentages of a specific proteoform within a background of another proteoform. A wide dynamic range is critical in proteoform assays as prior studies have shown tau isoforms can vary significantly in abundance ^29^. We serially diluted tau standard 2N4R-PKA (representing the tau proteoform 2N4R-pS214) into 0N3R, and performed seven detection cycles with five (anti-Tau13, anti-2N, anti-pS214, anti-4R, anti-Tau216) detection antibodies. Results indicate a linear dynamic range of 3.1, where the limit of quantitation was defined as the lowest point on the titration curve at which the signal to noise ratio is greater than 10 and the coefficient of variation is under 30%. (**Figure 3C**).

In summary, the tau proteoform assay, based upon the Iterative Mapping of Proteoforms method, quantifies tau proteoforms relevant to tauopathy, enabling reproducible detection of biologically meaningful variation using commercially available antibodies. Its analytical performance is linear across approximately three orders of magnitude for all tau isoforms, with a limit of detection of approximately 1000 ppm. This surpasses WB and offers insights beyond the reach of peptide-based approaches.

### Proteoform landscape of tau in models of neurodegenerative disease

Tau plays a central role in maintaining neuronal structure by stabilizing microtubules, particularly in axons. It has been extensively studied due to its pathological aggregation and hyperphosphorylation in FTD and AD, where it forms neurofibrillary tangles—a hallmark of neurodegeneration. Research studies to better understand tau’s physiological function and the molecular mechanisms driving its pathological transformation use a wide range of model systems, including iPSC-derived neurons, brain organoids, transgenic mice, and postmortem human brain tissue. Despite extensive study, the proteoform landscape of tau across model systems has yet to be established. To ensure that the assay had sample compatibility with the key sample types used in the field, we applied the tau proteoform assay to investigate what range of proteoforms were present, the characteristics of those proteoforms (e.g. were they dominantly singly modified species), and how those proteoforms differed across model systems. Our major goal in the below studies was to demonstrate that the assay could potentially be used to ask questions across a diversity of relevant model systems. As such, any biological findings introduced in the following sections should be treated as exploratory rather than definitive, particularly in light of the low number of biological replicates examined.

The model systems in the study included a mix of cell-derived models: 2D induced *NGN2*-induced neurons ^37^ (iNeurons) from four iPSC lines/donors and engineered 3D immuno-glial-neurovascular human multicellular integrated brain (miBrain) model with *APOE3* or *APOE4* genotypes to reflect AD disease risk from two lines, one donor at 8 and 16 weeks after hydrogel setup containing 9 and 17 week aged neurons ^38^. We additionally examined patient iPSC-derived cerebral cortical organoids comprised of approximately 16 subtypes of cortical neurons, astrocytes and progenitor cells, from 8 lines derived from 4 donors at 3 and 6 months carrying *MAPT* mutations (IVS10+16, V337M) and their isogenic controls (WT/WT) as a model for frontotemporal dementia (FTD) ^39,40^ (**Suppl. Table S4**), one hemisphere of brain tissue from a healthy hTau mouse (Mouse brain extract) and a frontal cortex tissue sample from a cognitively normal human (Human brain extract) (see **Methods** for more details about how samples were generated).

We compared the landscape of tau proteoform groups across the model systems, directly assessing their relative abundance. Samples from each model system were analyzed in at least three technical replicates. We note that this study does not attempt to estimate the biological variation of these model systems, but instead solely attempts to perform the first ever characterization of the proteoform landscape. A heat map of the normalized relative abundance of the tau proteoforms (mean across technical replicates) that were abundant at greater than a 0.1% level across the model systems is shown (**Figure 4A**). Models and proteoforms have been clustered by abundance (z-score normalized per-proteoform group). Each row corresponds to a different proteoform and each column represents a model system. The human brain and mouse brain samples had a lower number of phosphorylations. The stem-cell based models (iNeurons, Organoids and MiBrains) all showed higher extents of multiple phosphorylation. Notably, the cellular models were dominated by the 0N isoform, which is potentially most reflective of the highly phosphorylated tau species described in fetal tissue^41^. The mutant organoids at 3 months, in particular, showed the largest number of phosphorylated species.

**Figure 4:**
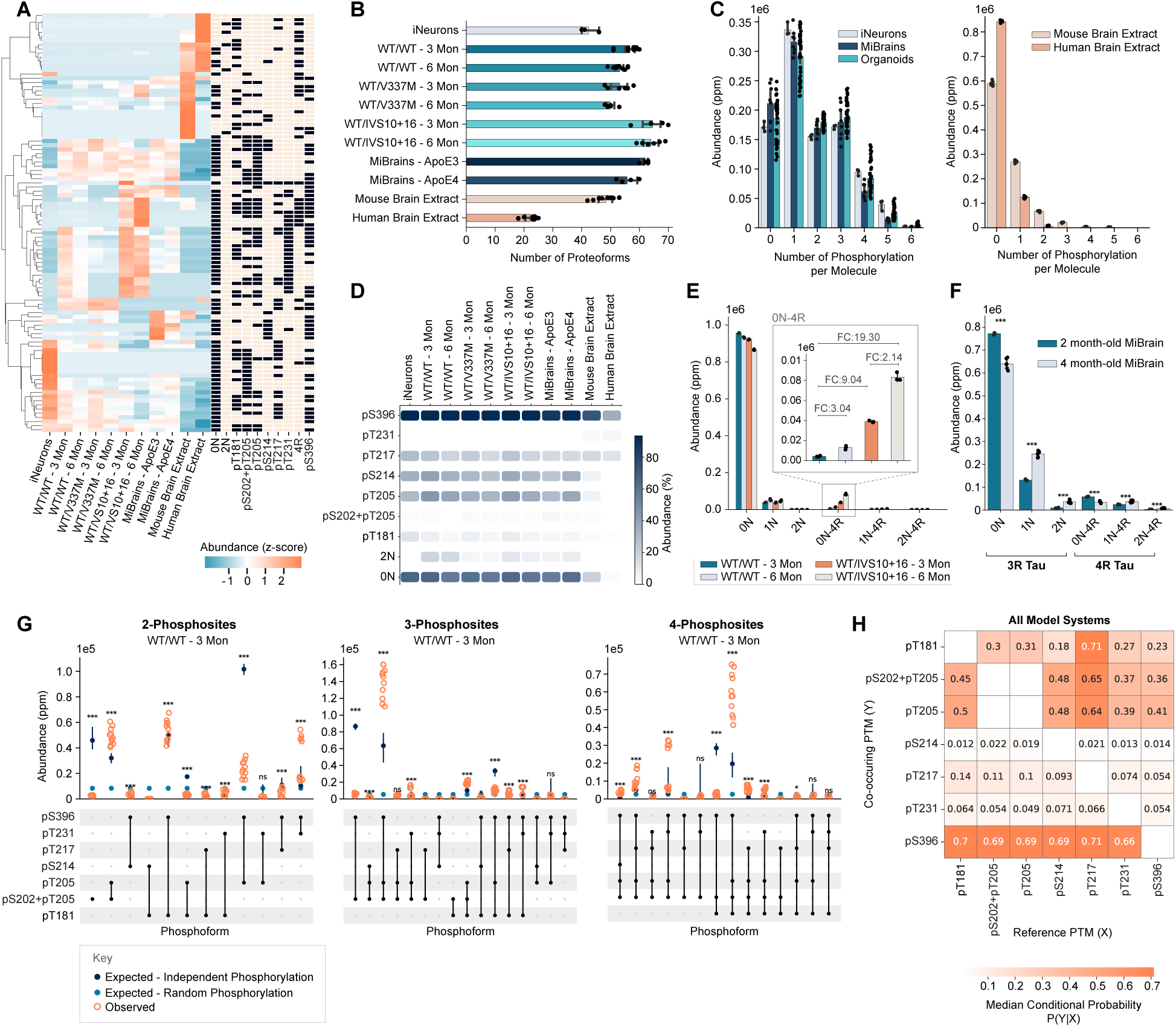
Examination of the landscape of tau proteoforms in models of neurodegenerative disease **Figure 4A:** Model systems (columns) and tau proteoforms (rows) are hierarchically clustered based on proteoform abundance. The blue-to-orange scale indicates relative abundance by z-score normalized per-proteoform of the mean of all replicates across each model system. The accompanying black/beige heatmap to the right denotes the presence (black) or absence (beige) of specific epitopes within each proteoform. **Figure 4B:** Number of proteoforms detected above 1000 ppm relative abundance for each of the tissues investigated. Data represent the RIPA buffer-soluble fraction of neuronal tissue as indicated. Number of phosphorylations are in Suppl. Table S5. For iNeurons n=3 technical replicates, for WT/WT-3Mon and 6Mon n = 12 (2 donors x 2 conditions x 3 technical replicates), for WT/V337M 3Mon and 6Mon n = 6 (1 donor x 2 conditions x 3 technical replicates), for WT/IVS10+16 3Mon and 6Mon n = 6 (1 donor x 2 conditions x 3 technical replicates), for MiBrains *APOE3* and *APOE4* n=8 (2 x 4 technical replicates), and for Human Brain Extracts and Mouse Brain Extracts n=9 technical replicates each. Error bars are standard deviation. **Figure 4C:** Number of tau protein molecules phosphorylated in human brain, mouse brain, miBrains, cortical organoids, and iNeurons. Data represent the average of at least two biological with 3 technical replicates with exception of the human brain extract (n=1). For iNeurons n=3 technical replicates, for Organoids n=24, for MiBrains n=8, and for Human Brain Extracts and Mouse Brain Extracts n=9. Error bars are standard deviation. **Figure 4D:** Pseudo-western showing the mean % occupancy for each tau epitope targeted in the assay across the replicates described in 4B. **Figure 4E:** Relative abundance (in ppm) of tau isoforms in maturing organoids at 3 and 6 months comparing IVS10+16 mutant (WT/IVS10+16) against isogenic controls (WT/WT). Representative result (donor GIH178C1) of two iPSC donor cell lines analyzed with an average of 8 organoids processed in two independent analyses. Cortical organoids details in Supplemental Table S4. Error bars are standard deviation. **Figure 4F:** Bar graph shows the relative abundance in ppm of the 3R and 4R tau protein in maturing miBrains of the *APOE4* genotype. Tau epitopes on the x-axis reflect the isoform specific probes used to assess full length tau protein isoforms (3R Tau putative). Noise level for the 3R tau was determined at 1474 ppm and at 485 ppm for the 2- and 4-month-old miBrains, respectively. For the 4R tau, corresponding noise levels for 2- and 4-month-old miBrains were at 1762 ppm and 702 ppm, respectively. Data shows four technical replicates of lysate from four miBrains at (57 days) and the average of 6 miBrains (112 days) processed independently. **Figure 4G:** Abundances are calculated independent of tau isoform and defined as phosphoforms. Orange dots represent observed phosphoform abundance, while dark blue and light blue dots indicate expected abundance based on models of independent and random phosphorylation, respectively. Statistical significance between observed and expected abundance levels from the independent phosphorylation model is shown (*p<0.05, **p<0.01, ***p<0.001). Error bars are standard deviation. N=12. **Figure 4H:** Co-occurrence of tau phosphorylation. Here we quantify across all model systems how often a given site is phosphorylated (y-axis) assuming the presence of a phosphorylation on another site (x-axis). The pattern of co-occurrence is distinctively non-uniform suggesting a preference for multiple phosphorylation.

Given the quantitative nature of the tau proteoform assay, we compared proteoform counts, uniqueness, and phosphosite occupancy across the diverse set of model systems and tissue samples analyzed. We define a proteoform group as observed if it appears in triplicate in at least one model system at an abundance >0.1% of all full-length tau species (non-proteolytically cleaved). We observed a broad range in the number of detected tau proteoforms, with most samples derived from stem-cell based models yielding between 50–65 distinct proteoform groups and only 23 tau proteoform groups observed in the healthy human brain sample (**Figure 4B**). Across all sample types, we observed 130 unique full-length (non-proteolytically cleaved) tau proteoform groups out of the 768 targeted by our assay (Supp. Table 5). We quantified the number of proteoforms shared across the models and found that out of all tau proteoforms detected, 34 proteoforms were found in only one model system, with fewer proteoforms shared amongst two or more model systems (**Suppl. Fig. 4A**). Overall, there was significant variation across the diverse models, with iNeurons appearing to be a large outlier relative to the organoid models (**Suppl. Fig. 5A**). Seven of the eleven proteoforms common to all model systems were phosphorylated, while all proteoforms found in only one model system carried phosphorylation. When comparing the proportion of phosphorylated to unphosphorylated tau molecules across all model systems, we observed the highest modification rates in WT/IVS10-16 cortical organoids (82%), followed by miBrain *APOE4* genotype (77%), WT/V337M mutant cortical organoids (76%), miBrain *APOE3* genotype (72%) and lower modification rates for the hTau mouse (37%) and human brain extract (13%) (Supp. Fig. 4B).

Next, we investigated the phosphosite occupancy across all model systems and found that the number of phosphorylations per tau molecule was much lower in animal model systems than in cellular model systems, where up to six co-occurring modifications per tau molecule were detected (**Figure 4C**). In contrast, tau proteoforms from mouse and human brain tissues displayed up to three concurrent phosphorylation events at significantly lower levels. Phosphorylation site occupancy analysis revealed pS396 as the most occupied epitope in cellular systems (60%), while pT181 showed the highest occupancy in animal models (25%). Notably, pT205 and pS202+pT205 were each occupied at ∼40% in cellular systems, exceeding their occupancy in the animal model (**Figure 4D**, **Supp. Fig. S4C**).

### Age-dependent changes in the tau isoform landscape

Tau splicing is regulated developmentally and only the shortest isoform (0N3R) is present in the fetal brain and maturing cellular models. This isoform may contribute to cytoskeletal plasticity during early neural development by enabling dynamic cytoskeletal adjustments needed in immature neurons ^42,27^. Adult tau isoforms containing 4R, on the other hand, are more efficient at promoting microtubule assembly^43^. The inclusion of three isoform-specific probes targeting 0N, 2N, and 4R tau allows our assay to resolve isoform-dependent biological differences. To assure that we can detect and quantify known isoform changes in tissue, we applied the tau proteoform assay to iPSC-derived organoid models containing the *MAPT* mutation ^40^ (IVS10+16), which alters *MAPT* splicing and results in increased levels of exon 10-containing mRNA (4R). This mutation skews the normally balanced 3R/4R tau ratio of 1:1 in the adult human brain towards 4R but does not affect tau’s primary sequence ^44^. Nevertheless, this imbalance leads to FTD-related neurodegeneration^44^. Comparing wild-type organoids (WT/WT) at 3 and 6 months of age, we detected an age-dependent increase (3.0-fold) in the mature 4R tau isoform with a similar, age-dependent increase (2.1-fold) for the mutant organoids (**Figure 4E**). Compared to wild-type organoids (WT/WT) at 3 months of age, organoids with the heterozygous *MAPT* mutation (WT/IVS10+16) at the same age had 9.04-fold more 4R tau isoform, increasing to 19.3-fold for the 6-month-old mutant organoids.

Similarly, we determined the age-dependent tau isoform ontogenesis when we measured tau isoforms isolated from 2- and 4-month-old miBrains. The engineered 3D immuno-glial-neurovascular human multicellular integrated brain (miBrain) model combines the enhanced neuronal maturation of organoids with the co-assembly of all six major central nervous system (CNS) cell classes into an engineered human brain mimic supporting the study of genetic risks factors like apolipoprotein E4 (*APOE4*) in sporadic AD. Here, the ability to detect full length tau protein allowed us to capture all six isoforms and determine the age-dependent ontogenesis of tau with relative abundances shifting from primarily 0N isoforms to the longer 1N and 2N forms (**Figure 4F**). The ability to directly probe the 4R isoform revealed the presence of the most mature 2N4R tau isoform at 7223 ppm, above the noise level of 1762 ppm. During aging of the miBrains, we observed a marked shift in tau isoform abundance from predominantly 0N3R to more mature isoforms, including 1N3R and 2N3R/4R. This transition from shorter to longer tau isoforms had been previously shown in humans ^43^. Comparing the relative isoform abundance of miBrains to human brain, the increased abundance of 2N4R in miBrains at 4 months of maturation more closely aligns with the adult human brain ^29^.

In summary, we recapitulated age-dependent changes in tau isoforms in a *MAPT* mutant model system of tauopathy. Furthermore, our assay captured isoform-specific changes in miBrain models. These findings show that our assay captures known biological phenotypes while offering greater specificity and broader tau proteoform coverage than traditional methods.

### Co-occurrence of multiple phosphorylation across model systems

Multisite phosphorylation may alter tau’s conformation and interactions with other proteins. Early stages of abnormal tau processing are believed to be characterized by sequential appearance of pT181 and pT231 followed by pS202 and pT205 then pS214 and pT217 ^45^. However, it has not been possible to directly observe if these modifications co-occur on individual molecules or instead are simply spread throughout many molecules. In general, co-occurrence of phosphorylation at one site is influenced by its own relative abundance and its preferred association with another site. Additionally, hyperphosphorylation of tau has often been mentioned as a hallmark of disease progression. However, it is unclear if hyperphosphorylation implies a greater percentage of molecules with any arbitrary phosphorylation, a greater percentage of molecules with a specific phosphorylation, or instead a greater percentage of tau molecules with multiple specific phosphorylations.

As part of demonstrating the potential applications of the tau proteoform assay, we investigated if it were possible to examine if tau has a specific pattern of multiple phosphorylation or if, instead, we simply see a relatively random multiplicity. With 7 anti-phosphoantibodies, there are 21 possible patterns of double phosphorylation and 35 patterns of triple or quadruple phosphorylation independent of isoform. While we observed 18 out of 21 patterns of double phosphorylation across the total set of samples, we only observed 16 of the 35 possible patterns of triple phosphorylation (**Supp. Table S6**). To assess whether multiple phosphorylations follow a random or preferential path, we calculated how abundant phosphoforms might be expected from a random phosphorylation path. We explored two random models, one in which modification was uniform random, and a site-dependent model that maintained the observed total extent of modification at each site. In **Figure 4G** we highlight this difference in observed phosphorylation patterns (orange) vs those expected at random by either a uniform random model (light blue) or a site independent phosphorylation model (dark blue). The assessment is made irrespective of the isoform in which the modification occurs and defined as phosphoform. Some phosphoforms with double, triple, and quadruple phosphorylation include sites T205, T217, and pT231 which exhibited greater than expected abundance across model systems, whereas many others showed abundances that agreed with an independent phosphorylation model (**Suppl. Fig. S4D**). For example, for the mutant organoid (IVS10+16) FTD model, we observed 3-fold higher abundance of triple (pT205-pT231-pS396) and quadruple (pT181-pT205-pT231-pTS396) phosphoforms than expected in an independent phosphorylation model (**Figure 4G**).

We next assessed the data for evidence of interactions between sites – where the phosphorylation status of one site influences the phosphorylation of another on the same molecule. We first looked at co-occurrence of phosphosite pairs. Some phosphosites co-occur with other phosphosites at higher rates than others. For example, pS396 occurs with each other phosphosite 66-71% of the time (**Figure 4H**). This is partially due to the extensive pS396 phosphorylation. We also assessed whether some phosphosites were more likely to be observed with another phosphosite rather than without it. Some phosphosite pairs appeared to have stronger interactions than others **(Figure 4H, Supp. Fig. S4E**). For example, the presence of pT181 increases the probability of observing pT217 8-fold (**Suppl. Fig. S4E**). Other phosphosites also notably increase the probability of observing pT217 including pS202+pT205 (5.2-fold), pT205 (4.3-fold), and pS214 (3.3-fold). Generally, pT217 appears to be the site most tightly coupled to the phosphorylation status of other sites on the same molecule. Conversely, pS396 appears to be the least tightly coupled to the status of the other sites on the same molecule, perhaps because it is physically further away, and outside of the proline-rich region of tau. This is despite pS396 being the highest abundance phosphosite observed in the data. Phosphosite dependency trends appear largely conserved across model systems with some evidence of higher codependency in brain extracts (data not shown). The dependence of pT217 on modification of other sites, without those sites having as substantial a dependence on pT217, suggests that pT217 may be a later modification, or suggest the activity of a phosphatase.

Collectively, the results highlight a complex and evolving landscape of tau phosphoforms, where the co-occurrence of distinct phosphorylation events suggests the existence of preferential and coordinated modification pathways and the conservation of co-occurrence across model systems implies conservation of said pathways. Notably, some of these co-occurring phosphorylations span the entire length of the protein—from the N- to the C-terminus—underscoring the necessity of a methodology capable of resolving combinatorial PTM patterns across the full-length, intact tau molecule.

### Tau proteoform assay compatibility with normal and cognitively impaired brain samples

To validate that the IMaP method was compatible with human clinical samples we analyzed a small cohort of 7 human samples that included aged asymptomatic patients and patients with Alzheimer’s Disease and Related Dementias (ADRD) (**Figures 5A, Supp.** Fig. 5**, Supp. Table S7**), Previous studies characterizing tau PTMs in individuals with AD have revealed considerable heterogeneity ^29^. We sought to determine whether our tau proteoform assay could measure human clinical tissue samples and if tau proteoforms could potentially distinguish between cognitively impaired cases and cognitively normal controls. We measured tau proteoforms from brain samples of a small cohort of five cognitively impaired patients with ADRD and two cognitively normal subjects. Clustering of impaired and normal samples revealed three distinct groups, with the normal samples clustering together. Nine proteoforms differentiated the normal samples from the ADRD samples. These nine proteoforms were significantly lower in abundance in ADRD samples (**Figure 5B**). These proteoforms contained phosphorylation at pT181, pT231, or both sites and were found on 0N3R, 0N4R, 1N3R, and 1N4R isoforms. While this is a relatively small sample size, these results suggest that tau proteoforms from brain tissue may differ between ADRD and normal.

**Figure 5:**
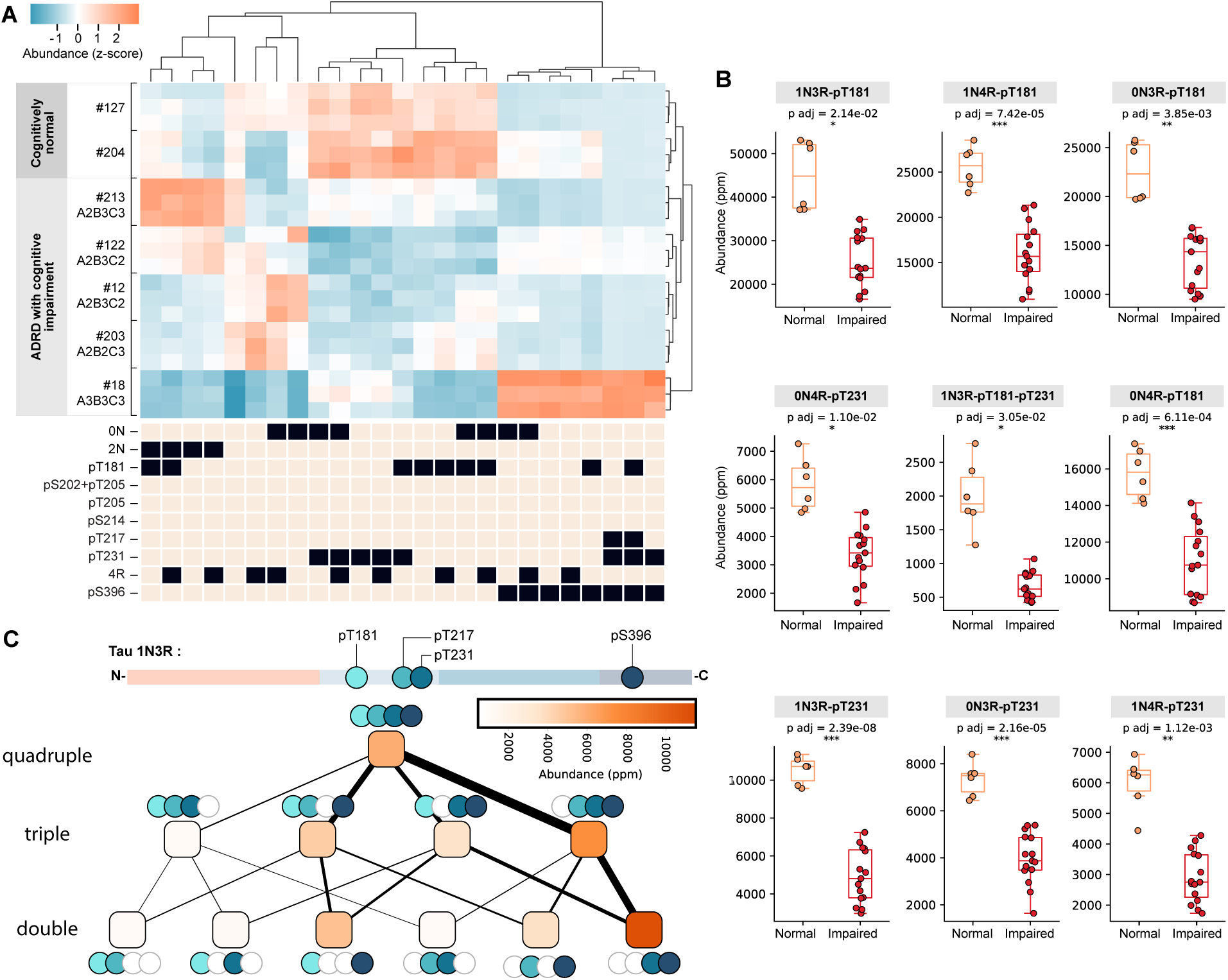
Application of tau proteoform assay to human samples **Figure 5A:** Tau proteoforms contrast normal from cognitively impaired brain samples. A heat map of the normalized relative abundance of the proteoforms of tau (mean across technical replicates) from samples of a small cohort with two cognitively normal patients (patient 127 & 204) and five cognitively impaired ADRD patients (18, 213, 122, 12, 203). Each patient is represented by three technical replicate measurements. Proteoforms plotted were at greater than 10% in abundance. Presence and severity of AD-related lesions for each patient is indicate in the lower panel following the ABC score. ABC = Thal, Braak, CERAD score for amyloid plaque deposition, tau protein tangles, and cortical neuritic plaques, respectively. **Figure 5B:** Relative abundance of multiple phosphorylations in human brain with two cognitively normal patients (patient 127 & 204) and five cognitively impaired ADRD patients. Tau proteoforms with multiple phosphorylations comparing cognitively normal to ADRD impaired. Star indicates p-values adjusted for multiple hypotheses using a Bonferroni correction. **Figure 5C:** Hyperphosphorylation of 1N3R in patient 18 at pT181, pT217, pT231, and pS396 potentially follows a preferential path from double to quadruple phosphorylation. Node fill color is the abundance of the proteoform as estimated by the mean of technical replicates. Edge weights correspond to estimated relative contribution of a given node to its parent nodes.

In addition, one patient with ADRD exhibited higher phosphorylation than the remaining patients. This patient also exhibited the most severe pathology, with an ABC score of 44 (A3, B3, C3), reflecting the highest levels of Aβ/amyloid plaques, neurofibrillary tangles, and neuritic plaques (each scored 0–3, with 3 indicating severe pathology)^46^. This patient harbored tau proteoforms with quadruply phosphorylated 1N3R tau at pT181-pT217-pT231-pS396. The doubly and triply phosphorylated forms of 1N3R that are a subset of the quadruply phosphorylated form suggest a regulated, rather than random, process driving these phosphorylation events (**Figure 5C)**. Three of four triply phosphorylated 1N3R proteoforms showed higher than expected abundance, whereas four of six doubly phosphorylated 1N3R proteoforms showed lower than expected abundance if phosphorylation events were randomly distributed (Supp. Fig. 6). These data suggest a preference for specific hyperphosphorylated tau proteoforms in the patient with the most severe ADRD.

## DISCUSSION

We have introduced a new method, Iterative Mapping of Proteoforms, for the large-scale analysis of intact single-molecule proteoforms and applied it to develop an assay for quantification of defined sets of proteoform groups of tau. As shown in **Figure 3** we observed a strong accuracy (R² = 0.985) across defined mixtures and median CVs below 5.5% across a wide range of reproducibility challenges. Currently, the greatest source of variation is in library preparation across operators. It is possible that further improvements in reproducibility could come from improvements in the robustness of the library preparation method and in improvements in training to further reduce operator-to-operator variation. Despite being a first demonstration of a new method, this CV is notably lower than those reported for existing, mature proteomics platforms ^47,48^. The assay resolved distinct proteoforms across greater than three orders of magnitude in dynamic range, providing a precise readout of relative stoichiometry of tau phosphorylation, and outperformed conventional methods like western blotting in distinguishing closely related proteoforms. In addition, for the first time, we reveal the molecular heterogeneity of tau proteoforms in model systems of neurodegenerative diseases iteratively mapping millions of single molecules to quantify the prevalence of distinct proteoforms.

Application of the tau proteoform assay allowed us to begin investigating several longstanding questions in the field of tau biology and proteoform analysis. First, we observe that tau molecules frequently harbor combinations of more than two phosphorylated sites, with certain proteoforms displaying up to six concurrent modifications. We additionally demonstrated that disease models do not just contain a greater percentage of phosphorylated tau molecules (82% for organoids vs 36% for healthy mouse brain and 13% for healthy human brain), but also that the number of phosphorylations-per-protein-molecule is larger (median 3/molecule in organoid vs 1.5/molecule for healthy human brain) reflecting the developmental or adult biology of the samples or their *in vivo* and *in vitro* origin, respectively.

Importantly, phosphorylations are not randomly distributed. We observed a non-uniform distribution of multi-phosphorylated tau species, with certain combinations of phosphorylation occurring disproportionately, suggesting an ordered and regulated process rather than stochastic modification. This is exemplified by the preferential co-occurrence of pT217 with pT181 with other co-occurrence of pT217 less frequently or not observed, implying temporal or hierarchical control in phospho-tau biogenesis. It is not possible to know whether this arises because of the action of kinases or phosphatases.

We further established that the extent of proteoform detection is limited not by assay sensitivity, but by biological diversity. Of the 768 possible tau proteoform groups detectable by our antibody panel, only a subset—ranging from 23 to 63, depending on the model—were actually observed. This suggests that tau modification is constrained by biological regulation, and not all theoretically possible proteoforms are realized in vivo. Previous work on histone revealed a similar extent of modification quantifying between 70-200 proteoforms for histone H4 and H3.2, respectively^,49^. Nonetheless, our approach dramatically expands the number of resolvable proteoforms compared to traditional methods.

Finally, our method and assay fill key gaps in the field. It enables (1) direct observation of site-specific co-occurring PTMs on individual molecules, (2) quantification of isoform-specific changes over time in human-relevant systems like organoids and miBrains, and (3) comparative assessment of model systems against human disease samples. Moreover, in at least one ADRD patient, we infer a plausible sequence of phosphorylation events leading to a distinct, highly-modified tau species, reinforcing the notion of ordered PTM accumulation in disease progression.

While Iterative Mapping of Proteoforms provides high-resolution quantification of proteoform groups, as demonstrated here with tau, several limitations should be considered. First, this study focused exclusively on full-length, RIPA-soluble tau, in contrast to most prior work that emphasizes sarkosyl-soluble and -insoluble fractions. This may particularly impact the interpretation of samples with a highly variable extent of truncation as our estimates of relative fold-change could be impacted by biological changes in total % of intact proteoforms. This can be addressed either through relating our estimates of proteoform relative abundance to all the measurements on the flow cell, all the measurements of tau of any length, or through using additional cycles for reagents that map truncations of tau explicitly.

Second, the targeted nature of this application of the method inherently limits its scope to known epitopes. Because Iterative Mapping of Proteoforms relies on predefined antibodies to detect specific isoforms and PTMs, it cannot identify novel sites or unanticipated modifications. This contrasts with discovery-based techniques like TDMS, which can uncover unexpected PTMs but lack single-molecule resolution and accurate stoichiometric co-occurrence information.

Third, this application of the method depends on antibodies with high specificity and compatibility with single-molecule detection. Cross-reactivity and inadequate epitope resolution—especially for adjacent or closely spaced modifications—can compromise accuracy. To mitigate this, we applied a dual-screening process for antibody validation and incorporated both null-lane controls and correction algorithms during data analysis, which effectively reduced non-specific signals to below 1%. Nonetheless, discriminating between closely spaced PTMs (e.g. neighboring phosphorylation sites) remains a fundamental challenge for antibody-based approaches. One possibility may also be to use highly non-specific affinity reagents, such as pan pTyr reagents in the assay to get a general gestalt of phosphorylation as a complement to the site-specific reagents used herein.

Fourth, caution is warranted when interpreting phosphorylation changes observed in this study, particularly in comparison to findings from single-epitope measurements. By design, single-epitope assays conducted on unenriched samples yield isoform-independent data, whereas the single-molecule proteoform analysis presented here captures isoform-dependent differences. Furthermore, absolute quantification of total tau relative to a multiply phosphorylated variant is challenging using both immunoassay and mass spectrometric methods. This distinction is especially relevant in the context of our small cohort study, where we observed a statistically significant reduction in singly and doubly phosphorylated tau proteoforms in impaired versus normal samples. This observation contrasts with the commonly held view that tau phosphorylation increases in disease (**Figure 5C**). Several factors may contribute to this apparent discrepancy including: the limited sample size, the presence of mixed ADRD pathologies and differences in protein extraction methods (RIPA vs. sarkosyl).

Finally, the current library preparation protocol requires lysine-based bioconjugation, which introduces random attachment points and molecular orientations on the array. While our results suggest that random orientation under denaturing conditions does not impede detection, future efforts to implement site-specific (e.g., N-terminal) labeling could further standardize molecule presentation and reduce potential sources of variability. Collectively, these limitations define the current boundaries of the tau proteoform assay and present opportunities for further refinement.

Iterative Mapping of Proteoforms establishes a new benchmark for high-resolution, single-molecule proteoform analysis, and several future directions hold promise for expanding its impact across neurodegeneration research and beyond. While protein sequencing technologies and TDMS offer broad discovery capabilities for detecting proteoform modifications, IMaP provides a scalable, targeted approach with unprecedented resolution of individual proteoforms beyond the reach of TDMS. The method is not limited to 12 antibodies and can be further expanded with additional probing cycles to include a wider array of PTMs. In stability studies (data not shown) we have demonstrated binding and removal of antibodies for over 150 cycles without loss of specific antibody binding or significant loss or damage of deposited proteins. Tau-specific modifications including acetylation, ubiquitination, and proteolytic truncations implicated in tau pathology are of high interest. Incorporating these dimensions will improve the granularity of tau proteoform characterization, particularly in disease progression and therapeutic response contexts.

With an observed median CV less than 5.5%, the reproducibility of the current assay now supports scaling to larger, biological studies and clinically annotated cohorts. Applying the tau proteoform assay to a broader range of brain samples—from cognitively healthy individuals to those across the full spectrum of AD and other tauopathies—may provide deep insight into the spatial and regional distribution of tau proteoforms. Leveraging pathologically characterized brain bank tissues could further link specific proteoform signatures to tau burden across multiple brain regions, generating maps of proteoform pathology that could inform disease staging and therapeutic targeting.

A critical next frontier is the adaptation of the assay for detection of tau proteoforms in biofluids such as cerebrospinal fluid (CSF) and blood, where concentrations are expected to be orders of magnitude lower than in tissue. Achieving single-molecule resolution in these matrices would enable the discovery of clinically relevant biomarkers and facilitate longitudinal studies to monitor the emergence and clearance of specific tau species over time. Such capabilities could support early diagnosis, stratification of disease subtypes, and assessment of drug efficacy in both preclinical and clinical settings.

Finally, the versatility of the method allows it to be readily adapted for other key proteins implicated in neurodegenerative disease—including α-synuclein, amyloid-β, and TDP-43—as well as proteoforms central to other human diseases such as cancer and autoimmune disorders. By applying the same principles, single-molecule Iterative Mapping of Proteoforms could help unravel the structural and functional heterogeneity of diverse protein populations, offering a transformative tool for proteoform-centric biology and precision medicine.

## MATERIALS & METHODS

### Screening and validation process for suitable antibodies

To assess the binding characteristics and specificity of the probe affinity reagents used for Iterative Mapping of Proteoforms we characterized the affinity reagents using a two-step process. We first assess the specificity of the antibodies to individual epitopes using control peptides (**Supplemental Table S1**). In a second step we confirm the specificity of the reagents (**Supplemental Table S3)** on the single molecule proteoform detection system to verify that those detection antibodies accurately recognized the specific epitopes of tau including phosphorylation and multiple phosphorylation sites.

As an example, to demonstrate that we accurately detect a mixture of three tau epitopes, we create phosphorylated peptides that represent tau epitopes of pT181, pS215, pS396 and immobilize them in a flow cell at a ratio of 1:3:1 to detect them with the corresponding antibodies. As negative control, corresponding non-phospho peptides or full length 2N4R proteins were used. Results from these specificity assessments using the negative control demonstrated that single molecule on-platform epitope recognition is qualitatively accurate and shows minimal off-target detection (cross-reactivity) with false positive detection below 1%.

Enrichment antibodies were selected for high affinity towards pan Tau proteins while still suitable for elution with cognate epitope peptides. They were able to deplete >90% Tau after overnight incubation with a protein mixture containing about 1 ng/ul Tau in ∼100 ul volume. The three antibodies used in this study recognize N-, middle, and C-terminus of Tau.

### Preparation of detection antibody conjugates

#### Modification of Anti-Tau Antibodies

Antibodies were acquired as carrier-free formulations in PBS pH 7.2-7.4 (**Supplementary Table S3**), or else buffer exchanged into PBS pH 7.4 using Amicon Ultra 100k MWCO Regenerated Cellulose Spin Filters. Modification is performed by first adjusting the pH of the antibody solution using 0.5M Sodium Phosphate (pH 8.0, 0.25 ml per of Antibody). A freshly prepared solution of sulfo-NHSLC-LC-Biotin is added to the Antibody solution in 20-fold molar excess and incubated (25C, shaking, protected from light) for 2 hours. Quenching of the modification reaction is performed for 0.5 hours (25C, shaking, dark) with a final concentration of L-Arginine (in the quench reaction) of 10mM. Modified antibodies are purified using desalting column (Zeba, Thermo Scientific). The extent of biotinylation is measured using MALDI Mass spectrometry

#### Conjugation of Anti-Tau Antibodies to fluorescent labels

Fluorescent labels containing up to four streptavidin groups are incubated with 2 molar equivalents of biotinylated anti-Tau antibody with equivalents calculated relative to the measured number of Streptavidin attached per nanoparticle. The conjugation proceeds for 2 hours at 25C, shaking in the dark. Ab-conjugated nanoparticles are purified from excess unconjugated antibody via HPLC-Size Exclusion Chromatography (SEC) using an Agilent 1260 Infinity II HPLC system equipped with DAD and fraction collector. The SEC column used is a Shodex OHpak SB-2006M (P/N: F6516017). Method for purification consists of isocratic flow at 2.5mL/min of 1x HPLC O.B. (200mM NaCl, 11mM MgCl_2_, 5mM Tris-HCl, 1mM EDTA) for 1.5 column volumes. Fractions corresponding to the nanoparticle peak are pooled and concentrated using Amicon Ultra 100k MWCO Regenerated Cellulose Spin Filters. Purified, concentrated sample is stored in DNA Low-Bind tubes at 4C.

#### Tau proteoform standards and mixtures

To assess the accuracy and performance of our platform to correctly identify and quantify specific tau proteoforms, we created four highly characterized tau protein standards and mixtures thereof. Tau proteoform standards, included a) unphosphorylated tau 0N4R isoform, b) the unphosphorylated 2N4R isoform, c) phosphorylated tau 0N4R isoform treated with ERK2 (multiple phosphorylation) and d) the tau 2N4R isoform treated with PKA (single phosphorylation at serine 214). We established the exact composition of four distinct tau proteoforms by LCMS assessing the site and percentage of phosphorylation of tau proteoform standards as described in the Supplemental Tables and Figures (Table S2).

Additionally, we purchased tau standards 0N4R (BioLegend 843001), 2N4R (R&D Systems (SP-495). The tau standards were used throughout the study as proteoform controls and used to prepare tau proteoform mixtures as indicated in the experimental details. As binding controls for the assay, we created DNA-origami nanostructure conjugates of the following tau proteoforms standards 2N4R, 0N4R, 0N3R, triply and quadruply phosphorylated peptides (see **Suppl. Table S1**), a short peptide as a null target, and standard mixtures of 0N4R:2N4R:2N4R-PKA tau proteoforms at different ratios as indicated in the experimental details. Before mTz modification, tau proteoform standards commercially purchased recombinant Tau proteins were first precipitated with four volumes acetone and resuspended in PBS, supplemented with 1% SDS.

#### Cells and Tissues

Postnatal-like iPSC-derived neurons (iNeurons) were generated using the *NGN2* protocol and cultured as previously as described ^50^. Cortical organoids from patient-derived iPSC using a guided differentiation protocol with dual SMAD and WNT inhibition in a 96-slitwell format, as previously described ^51^. 10 organoids at 3 months and 6 months maturity, harboring the original disease genes or reverted to the wild type by CRISPR techniques (**Supplemental Table S4**), were harvested as 3 vials (3-4 organoids each) for protein extraction and library preparation in parallel. ^38,5138,5238,5138,5238,5338,52^. 4-6 miBrains harboring apolipoprotein E3 or E4 genes (*APOE3/E4*), were harvested individually at days 57 and 112 after seeding and processed for protein extraction and proteoform analysis. hTau mouse brains at 3 months age used in Figures 4 and 5 were obtained from Quest Pharmaceutical Services (Newark, DE), from transgenic mice expressing a human tau derived from a human PAC, H1 haplotype, and with the murine Tau gene knocked out by a targeted disruption of exon1. The healthy human frontal cortex brain sample denoted Human brain extract as used in Figures 4 and 5 was obtained from Analytical Biological Services Inc. (New Castle, DE) from a postmortem Caucasian male donor died at the age of 66 with lung cancer without neurological disorder. A cohort of seven post-mortem human brain samples from the cortical region was obtained from the Mount Sinai Neuropathology Brain Bank and Research CoRE. These samples are used Figure 5. Donor information and disease stages are listed in the table **Supplementary Table S6**.

A detailed description of all the biological samples and their lysis/protein extraction procedures can be found in **Supplementary Information**.

#### Single-molecule Iterative Mapping of Proteoforms Methodology

The method we introduce here, Iterative <u>Ma</u>pping of <u>P</u>roteoforms (IMaP) consists of four steps that are detailed below and diagrammed in Figure 1. The first step is library preparation (detailed in **Single molecule library preparation**) which aims to take protein molecules from a sample and conjugate each molecule to a DNA-origami nanostructure.

In one instantiation of this workflow recombinant protein is used directly functionalized with an amine-reactive mTz click reagent (as described in *Functionalization of recombinant Tau proteins).* In another instantiation of this workflow protein is extracted from tissue lysate prior to functionalization (as described in *Functionalization of cell lysates)*. If needed, a sample can be enriched using protein-specific antibodies (detailed in Optional *Tau Enrichment).* Next, protein is conjugated to DNA-origami-based nanoparticles. When used alongside enrichment, functionalization can either occur using protein directly (detailed in *DNA nanostructure conjugation for un-enriched samples*) or on bead (detailed in *DNA nanostructure conjugation for enriched samples*). At no point is the protein digested. Next (Figure 1B), protein-nanoparticle conjugates are deposited into a lane of a custom microfluidics flow cell ^34^. Flow cell fabrication is described in Supplemental Methods in the section (*Flow cell preparation and assembly*). Single molecules are iteratively probed (as described in the section *Sample analysis*) in a massively parallel manner using a series of site-specific fluorescently labeled ^35^ antibodies (Figure 1C). Antibody labeling is described above in the section (**Preparation of detection antibody conjugates**).

Each iteration consists of a bind, rinse, image, and remove cycle. Specifically, a fluorescently labeled antibody probe is introduced into the flow cell and allowed to incubate. Next, unbound antibody is rinsed from the flow cell and the flow cell is optically imaged ^34^ to determine where the antibody has bound. Following imaging, bound antibodies are removed from the flow cell, preparing the flow cell for a subsequent iteration. As fluorescent images are acquired, positions of each single-molecule binding event are recorded. Collectively, probing of the single-molecule array produces a series of binary bind/no-bind outcome patterns (single molecule binding traces), which are used to determine the potential identity of the proteoform at each coordinate in the flow cell. Analysis of the images and sample quantification are described in the sections *Processing of Images* and *Proteoform Analysis* respectively.

### Single molecule library preparation

#### Functionalization of recombinant Tau proteins

Protein samples are resuspended in PBS, supplemented with 1% SDS and then treated with 100 uM methyltetrazine-PEG4-STP (4-Sulfo-2,3,5,6-tetrafluorophenyl) ester (mTz) (Vector labs, CCT-1399) (with 1% DMSO) at 25C for 3 hours on a thermomixer shaking at 850 rpm protected from light. The reaction was quenched with Dithiobutylamine (Sigma Aldrich 774405) at 5 mM for 1 hr at 25C, followed by iodoacetamide (Pierce A39271) at 10 mM for 15 min at 25C. The protein was precipitated by 9 volumes of absolute ethanol, washed once with ethanol at room temperature, air dried, then solubilized in (200mM NaCl, 15mM MgCl_2_, 5mM Tris-HCl, 1mM EDTA, 1% SDS), and quantified using the Pierce 660nm Protein Assay Reagent following manufacturer’s manual (Thermo Scientific).

#### DNA nanostructure conjugation for un-enriched samples

For recombinant proteins, or eluted proteins mTz modified proteins were incubated with DNA nanoparticles (at 50∼100 nM) bearing a single TCO (*trans-*cyclooctene) functional group^53^ at an approximate molar ratio of 5:1 in a buffer (200mM NaCl, 15mM MgCl_2_, 5mM Tris-HCl, 1mM EDTA) overnight (approximately 16 hours) at 25C protected from light. Spin filtration with a 100kDa MWCO cut-off Amicon Ultracel (Millipore Sigma UFC910008) was used to remove excess unconjugated proteins and exchange to a new buffer (5mM Tris-HCl pH 7.5, 200mM NaCl, 15mM MgCl_2_, 1mM EDTA, 0.05% CHAPS).

A typical reaction of about 20 ul was first diluted to 500 ul, concentrated to about 20 ul, and repeated three times in total. The purified Tau-conjugated nanoparticles were quantified with a Nanodrop instrument using the dsDNA settings.

#### Functionalization of cell lysates

Neuronal tissue/cell lysates and Expi293 lysates with recombinant Tau spiked-in were first precipitated with 4 volumes of pre-chilled (−20C) acetone. The pellets were briefly washed with cold acetone, air dried, then resuspended in PBS supplemented with 1% SDS. Lysates were quantified with Pierce 660 Assay, adjusted to 2 mg/ml, then treated with 200 uM methyltetrazine-PEG4-STP (4-Sulfo-2,3,5,6-tetrafluorophenyl) ester (mTz) with 2% DMSO at 25C for 3 hours on a thermomixer shaking at 850 rpm protected from light. The reaction was quenched with Dithiobutylamine (Sigma Aldrich 774405) at 5 mM for 1 hr at 25C, followed by iodoacetamine (Pierce A39271) at 10 mM for 15 min at 25C. The protein was precipitated by 9 volumes of absolute ethanol, washed once with ethanol at room temperature, air dried, then solubilized in PBS supplemented with 1% SDS.

#### Optional Tau Enrichment

When used with biologic samples, enrichment of tau is performed using a magnetic bead-based pulldown. Anti-Tau monoclonal antibodies targeting N-, middle, and C-terminus of Tau, optionally together with mouse IgG (Invitrogen 31903) as the negative control, are immobilized individually using Dynabeads™ Antibody Coupling Kit (Invitrogen 14311D) at 10 μg antibody per mg beads according to the manufacturer’s instructions. Anti-Tau beads were mixed in equimolar ratios prior to use. Modified lysates were diluted by 5 volumes of IP Buffer (50mM Tris-HCl pH 8.0, 150mM NaCl, 1% Triton X-100) supplemented with 1x Protease/Phosphatase Inhibitor cocktail (Cell Signaling Technology #5872), to bring final SDS to 0.16%, then mixed with antibody containing beads at an IgG:Tau ratio of around 15∼50:1 at room temperature overnight (approx. 16hrs), on a tube rotisserie. The beads are then extensively washed with PBS and IP Buffer 2∼3 times each,

#### DNA nanostructure conjugation for enriched samples

While still on the beads, enriched tau is conjugated with 2.5 molar excess of DNA nanoparticles (relative to the estimated Tau input) bearing a single TCO (*trans-* cyclooctene) functional group in a buffer (5mM Tris-HCl pH7.5, 200mM NaCl, 15mM MgCl_2_, 1mM EDTA) overnight at room temperature on a tube rotisserie. Beads were washed three times with buffer (5mM Tris-HCl pH 7.5, 200mM NaCl, 15mM MgCl_2_, 1mM EDTA, 0.05% CHAPs), and eluted with the same buffer supplemented with synthetic peptides encompassing the epitopes of each of the Tau antibodies used at room temperature for 90 minutes on a tube rotisserie. Tau-conjugated nanoparticles were quantitated based on 488 florescence if the nanoparticles were labeled, or by Quant-iT PicoGreen dsDNA Reagent (Fisher Scientific P7581) following manufacture suggested procedure, using the unconjugated nanoparticles as standards, aliquoted and stored at −80C before use.

### Sample analysis

#### Experimental Set-up and Imaging

Flow cells, running buffer (50mM HEPES pH7.4 + 120mM NaCl+5mM KCl + 10mM MgCl2+ 0.1% Tween-20 + 0.5% Procilin 150+ 10mM Sodium Sulfite +10mM Sodium L-Ascorbate), remove buffer (5.2 M Guanidine HCl, 20 mM MgCl2, 20 mM Sodium Acetate, pH 4.30 +/- 0.10), DNA nanoparticle conjugated Tau molecules suspended in a buffer comprising 5 mM Tris, pH 8.0, 120 mM NaCl, 12.5 mM MgCl2, 1 mM EDTA, 0.05% CHAPS and affinity reagents diluted in 10 mM HEPES, pH 7.4 120 mM NaCl, 12.5 mM MgCl2, 5 mM KCl, 0.1% Tween-20, 1% Pluronic F-127, 1 mg/mL ssDNA, 1% BSA, 0.1% ProClin-150 were all loaded onto the instrument prior to assay initiation. For the duration of the assay, the running and remove buffers were held at room temperature (∼21° C) while labeled probes were maintained at 15° C and the flow cell surface was maintained at 23° C. The assay was performed on the instrument in an automated fashion involving sequential steps. In brief, the flow cell surface was primed with running buffer, followed by injection of the DNA nanoparticle-conjugated Tau proteins. Deposition onto the flow cell surface was mediated by the annealing of complementary oligonucleotides between the nanoparticle and pre-immobilized DBCO-oligonucleotides on the flow cell surface. After a 30-minute incubation, the unbound material was washed away from the flow cell using running buffer. Subsequently, sequential probing was carried out using fluorescently labeled antibodies targeting distinct epitopes. Each probe was incubated for 17 minutes, followed by a wash step with running buffer to remove unbound probes. A 17-minute incubation with remove buffer was then used to dissociate the bound probes, resetting the surface for the next cycle. This process was repeated for 36 total cycles, covering 12 unique probes.

Images of the flow cell surface were acquired at key checkpoints of each cycle using a custom-built high-resolution imaging system. This system included an Olympus UPLXAPO20X objective lens, Thorlabs TTL180-A 180mm focal length tube lens and dual Teledyne Dalsa Linea HS4K TDI line scan cameras. Excitation was provided by 488nm and 638nm lasers. With an assortment of dichroic and bandpass filters, multicolor imaging was enabled. Autofocus was maintained using a “Dover DOF-5” objective focusing stage integrated with a WDI ATF4 autofocus module. A high precision Dover SmartStage linear stage was used for positioning the flow cells under the objective lens and enabling continuous scanning of the flow cell surface throughout multiple cycles. A custom designed flow cell holder was mounted on the Dover SmartStage to maintain flow cell position and temperature stability of the flow cells under the microscope during imaging. Images of the flow cell surface were acquired after protein conjugated DNA nanoparticle deposition, following each probe wash step and after every remove buffer incubation step. These image stacks were then processed to perform alignment, spot detection and downstream data extraction for quantitation and analysis.

### Data analysis

#### Processing of Images

The input images were processed as previously described ^34^. Briefly, patterned array and single pad boundaries were determined using subarray fluorescent patterns excited by the 488 nm laser. The mapping between 488 nm and 647 nm laser acquired images uses subarray locations in the 488 nm scan and fiducial beads in the 647nm scans. A calibration flow cell is used to allow mapping of locations directly from 488 nm images to 647 nm images. A reference base scan is taken for both the 488 nm and 647 nm images. The 647 nm-reference scan is used to align all subsequent 647 nm images. Each image was normalized against the background. Background intensity estimates were calculated by tiling the image and computing the 5^th^ percentile for each tile. Then, the tiles are used to solve for a 2-dimensional paraboloid. The image is then normalized against this smooth paraboloid. The primary feature score was computed for each candidate object by summing the logarithm of 5 pixels forming a 3×3 t-shape centered on each landing pad in the background normalized image.

#### Proteoform Analysis

The Iterative Mapping of Proteoforms algorithm analyzes a series of antibody binding measurements acquired on a sample with unknown proteoforms and determines the abundance of each epitope in the sample. The algorithm uses an expectation-maximization approach to estimate the probability of each proteoform in the sample. The inputs to this algorithm are the series of antibody binding measurements and the rates at which the antibodies bind to their target epitopes. The model employs a Poisson binomial distribution, where the likelihood that the proteoform contains a given epitope is determined by the antibody binding rates. The output of the model is partial counts corresponding to the probability that each candidate proteoform produced the observed series of binding measurements. The number of candidate proteoforms is 2*^m^*, where *m* represents the number of antibodies used in the experiment. To account for sample-to-sample and run-to-run variation, the algorithm also estimates the binding rates for each antibody within each sample. We use control proteoforms to determine the affinity of labeled antibody binding in the presence and absence of the target epitope, and set a prior on antibody binding rates based on these control proteoforms. A uniform prior is set on proteoform abundance estimates. To account for potential protein degradation during sample preparation, we discard any landing pads that do not contain at least one positive binding measurement for an antibody that targets the N-terminal region of Tau and at least one positive binding measurement for an antibody that targets the C-terminal region of Tau. This ensures that we analyze landing pads that contain full length Tau. A two sided t-test was used to assess the significance of differences in abundance between proteoforms across samples. In the case of the cognitively impaired and non-cognitively impaired cohorts, p-values were adjusted to correct for multiple hypotheses using a Bonferroni correction.

#### Expected Multiply Phosphorylated Tau Molecules

Two models for expected phosphorylation abundance were constructed. A “Random Phosphorylation” model was built by assuming that all epitopes were equally likely to be phosphorylated. An “Independent Phosphorylation” model was built by assuming that phosphorylation events occur independent from phosphorylation elsewhere on the molecule. This process was modeled as a joint Bernoulli model, such that the probability of observing each combination of post translational modifications is equal to the product of the probabilities of the presence and absence of each individual phosphorylation event. In the case of Supplementary Figure S4D, isoform information was ignored. In Supp. Fig. S5, isoform information was included such that the expected phosphorylation values were based on the singly phosphorylated occupancies of the 1N3R isoform.

#### Co-Occurrence of Tau Phosphorylation

To assess whether post translational modifications were co-occurring at a higher than expected rate, we calculated the likelihood ratio comparing two conditional probabilities. We compared the rate at which post translational modification Y occurs given post translational modification X to the rate at which post translational modification Y occurs when post translational modification X is not present. Notably, this analysis ignored isoform information.

## Consortia

The members of NAUTILUS BIOTECHNOLOGY are Jamie E. Anderson, Fletcher Bain, Filip Bartnicki, Issa Beekun, Giovanni Bellesia, Rajat Bhatnagar, Keith Bjornson, Phillip S. Brereton, Stephen Brown, Vivekananda Budamagunta, Kaitlyn Burke, Chieh-Li Chen, Kevin Chen, Priscilla L. Chen, Krista Cosert, Danya Cruver, Michael A Darcy, Wade Dugdale, Ahana Dutta, Jarrett D. Egertson, Michael E. Flaster, Tyler J. Ford, Keith Frankie, Taryn E. Gillies, Keith Gneshin, Hamid R. Golnabi, Sam Grillo, Rob Grothe, Cynthia Y. Guerrero, Pengyu Hao, JSRoss Hartley, Brooks D. Hoehn, Jr., Huber Martin, Andreas FR Huhmer, Christina E. Inman, James Joly, Kevin M. Jin, Maryam Jouzi, Kara Juneau, Greg Kapp, Kota Kaneshige, Stanislav Katsyuk, Eun Kim, Rajesh Kota, Jamie Rose Kuhar, Jonathan B. Leano, Darrel Lee, Winnie Leung, Cara Li, Bryant Lowry, Parag Mallick, Mara Manskie, Carlos F Martinez, Kevin McVey, Jonathan C. Menger, Denise A. Miller, Eva Monsen, Anna Mowry, Grant Napier, Nicholas Nelson, Nicole Newland, Maureen R. Newman, Brittany Nortman, Adedeji Ogunyemi, Jayetha Panakkadan, Sujal Patel, Rukshan T. Perera, Nguyen Pham, Keshav Prabhu, Maximilian O Press, Hongji Qian, Torri E. Rinker, Adam J. Roberts, Julia K. Robinson, Aimee A. Sanford, Ali Najafi Sohi, Subra Sankar, Parag Shekher, Genee Victoria Skulich, Irene Tang Sparck, Isaac B. Sprague, Noah Steiner, Robert Stephenson, Juanfeng Sun, Kentaro Suzuki, Scott M. Tabakman, Steven J Tan, Nicholas Tardif, Sonal S. Tonapi, Jasmine Trinh, Tuan Trinh, Maria Villancio-Wolter, Jacinto Villanueva, Zhiqiang Wang, Lisen Wang, Gwen Weld, Sheri K. Wilcox, Katherine Winters, Feng Yan, Yvonne Yeung, Zhengjian Zhang, All members are affiliated with Nautilus Biotechnology, San Carlos, CA 94070.

## Author contributions

VB, SG, MJ, and JJ designed and executed the experiments on the Nautilus platform, RW, BN, and ZZ designed the Tau enrichment method, developed the sample preparation methods, developed the single molecule library preparation methods, and prepared the single molecule libraries for analysis. ST prepared peptide and proteoform standards. PB and JS prepared affinity reagents. JJ performed the data analysis. GK, KJ, PM conceptualized experiments and contributed to the manuscript. TB, SL, NeuraCell prepared and performed the organoid experiments. ST contributed to the manuscript. AR contributed to experimental design and strategy as well as research management, particularly with regards to recombinant tau strategy, choice of constructs and design of phosphorylation. LR contributed to research execution and materials in particularly with regards to expression and purification of phosphorylated tau. JL contributed to experimental design and materials particularly in regards to preparation of iNeuron cells. TJW contributed to research execution and materials in particularly with regards to expression and purification of phosphorylated tau. DA performed the mass spectrometry analysis and phosphomapping of pTau. ERS generated, cultured, and prepared multicellular integrated brains for protein analysis. JES Prepared multicellular integrated brains, performed complementary staining confirming tau presence and performed protein extraction on human brain samples. ALFJ generated, cultured, and prepared multicellular integrated brains and performed complementary stainings confirming tau presence. JB conceptualized experiments and supervised experiments. AFH, BN, CEI, KJ, MEF, PM, RG, SKW, VB, ZZ contributed to conceptualization. ANS, BDH, BN, GB, HQ, JDE, JKR, JT, JV, KJ, MEF, MJ, MRN, PI, PM, RG, RK, RTP, SG, SJT, SKW, SMT, VB, ZZ contributed to methodology. AJR, CC, DAM, DL, EM, HRG, KC, KM, KW, LW, MEF, MM, MV, NT, RB, RG, RH, SB, SK, WL contributed to software. AO, BN, JS, KJ, KM, KP, KW, MEF, MJ, MV, PSB, RG, RTP, SJT, SST, TER, VB, ZW, ZZ contributed to validation. FB, JDE, JS, KM, MJ, RG, ZZ contributed to formal analysis. AD, AO, BDH, BN, FB, FY, JBL, JS, JT, KB, LW, MAD, MEF, MJ, MV, NS, PI, RG, RTP, SG, SG, SMT, TT, VB, ZZ contributed to investigation. AAS, AD, BL, BN, CYG, DL, EK, FB, FY, GVS, JBL, JCM, JV, KB, KC, KF, KK, KM, KMJ, KS, MAD, MOP, NS, PSB, RS, TER, WD, YY, ZZ contributed to resources. AO, CL, FB, FY, KC, KM, MJ, MOP contributed to data curation. AFH, CEI, JJ, PM, VB, ZZ contributed to writing the original draft of the manuscript. AFH, DCB, BN, JDE, JJ, JRK, KJ, KS, ST, PM, TJF, VB, ZZ contributed to reviewing and editing the manuscript. JDE, JJ and PM contributed to visualization. FY, GW, HM, JRK, JV, KJ, MJ, PM, SKW, SMT, SP, ZZ contributed to supervision. DL, GW, JCM, JEA, MAD, PM, SKW, SMT, SP, YY, ZZ contributed to project administration. AM, PM, SP contributed to funding acquisition.

## Acknowledgements

*MAPT* Human induced pluripotent stem cells lines (**Supplemental Table S4**) were provided through the generous support of the Tau Consortium and the Rainwater Charitable Foundation and are available at https://www.neuralsci.org/tau. We are indebted to the NSCI NeuraCell core facility for organoid production and technical support.

We additionally acknowledge funding from The Regenerative Research Foundation (NSCI investigators); The Rainwater Charitable Foundation and the Tau Consortium and Cure PSP (ST, TB, DCB); NIH: RF1NS123568 (ST, DCB), 1RF1NS142335 (ST, TB). This work was funded in part through support to JWB from NASA (contract #80ARC022CA004), the NIH/NIA R01AG089533, and The CureAlz Fund.

## Additional information

Supplementary Information is available for this paper. Correspondence and requests for materials should be addressed to

## Supplementary Information

### Supplementary Materials and Methods

#### Recombinant protein production & characterization

Recombinant Tau proteins were expressed and purified following a protocol similar to Baghorn et al.^1^. Briefly, genes encoding for the 2N4R and 0N4R Tau isoforms were codon optimized, synthesized, cloned into a modified pET15 vector and transformed into BL21 (DE3) competent cells (Agilent, 230132); 100 µL of the reaction was plated onto LB plates overnight at 37°C. A single colony was used to inoculate a starter culture overnight; resulting culture was used to inoculate 500 mL of LB medium ampicillin medium with 5 ml; cultures were grown in UltraYield flask (Thompson, 931136-B) until OD_600nm_ reached ∼0.8 unit and induced with 0.5 mM isopropyl β-d-1-thiogalactopyranoside at 30°C for 3h and pelleted. Cell pellets were suspended in buffer A (25 mM HEPES pH 7.2, 50 mM NaCl, 1 mM PMSF, 0.5 mM TCEP, and cOmplete™ EDTA-free Protease Inhibitor Cocktail, 5 mM MgCl_2_ and benzonase (prepared in-house). The suspension was homogenized and passed through a microfluidizer twice to break the cells open. The lysed suspension was boiled for 20 min and spun down at 40000 rpm for 40 minutes. The supernatant was loaded onto a 5 mL pre-packed SP HP column (Cytiva, 17115201) equilibrated with Buffer B (25 mM HEPES pH 7.2, 50 mM NaCl and 0.5 mM TCEP). The column was washed with 10CV of Buffer B and eluted with a gradient of Buffer C (25 mM HEPES pH 7.2, 1 M NaCl and 0.5 mM TCEP, 0-50% of Buffer C over 30CV). Fractions containing protein were pooled, concentrated down and loaded onto a HiLoad 16/60 Superdex 200 pg column pre-equilibrated with PBS pH 7.2 with 0.5 mM TCEP. Protein concentration was determined via NanoDrop using the Protein A_280nm_ method.

The DNA sequence encoding for ERK2 kinase domain (Met1-Ser360) was cloned in a derived pET15b vector with a N-terminal TEV cleavable His-tag (MHHHHHHGENLYFQGS), transformed into BL21 (DE3) competent cells (Agilent, 230132) and grown as described above except that IPTG induction was allowed overnight at 16°C. Pellets were resuspended in ∼ 100 mL BugBuster (Millipore, 70921) supplemented with benzonase, lysozyme (purified in-house), cOmplete™ EDTA-free Protease Inhibitor Cocktail, 20 mM imidazole, and 1 mM TCEP for 1h and purified following Chen et al., (“Clearing the Path to Rapid High-Quality Protein Purification”, manuscript in preparation). Briefly, Ni-NTA magnetic beads were washed with Ni Buffer (50 mM Tris pH 8, 300 mM NaCl, 10% glycerol and 1 mM TCEP) supplemented with 20 mM imidazole and added to the lysate. The lysate was incubated at room temperature for ∼30 min on an orbital shaker. The beads were collected using a magnetic block and magnetic wand, washed with 5 x ∼25 mL of Ni Wash Buffer using a magnetic rack and eluted using Ni Buffer supplemented with 300 mM imidazole. The eluted protein fraction was incubated overnight with two aliquots of His-TEV protease (prepared in-house) and dialyzed against 2 L of Ni Buffer at 4°C. The digested protein was run over a 2 mL Ni gravity flow column, washed with Ni Buffer with 20 mM imidazole and eluted with 300 mM imidazole. Flowthrough and Wash fractions were collected, diluted to ∼400 mL in MonoQ Buffer A (20 mM Tris pH 8.5, 5% glycerol and 1 mM TCEP) and loaded onto the MonoQ 10/100 (Cytiva, 17516701). The column was washed with 20 CV MonoQ Buffer A and the protein eluted with a gradient to 50% MonoQ Buffer A supplemented with 1 M NaCl over 30 CV. Liquid Chromatography-Mass Spectrometry (LC-MS) analysis confirmed this protein was not phosphorylated; protein was concentrated down and loaded onto a S200 26/60 pre-equilibrated with SEC Buffer (20 mM Tris pH 7.5, 5% glycerol, 150 mM NaCl and 1 mM TCEP). Fractions were pooled, diluted to 0.1 mg/mL in SEC buffer. 0.02 mg/mL of a phosphomimetic constitutively active full length human MEK1 (Pro2-Val393) (mutations at Ser218Glu and Ser222Glu, prepared in house) and 5 mM MgCl_2_ were added. 1 mM ATP was added to initiate phosphorylation reaction. The reaction was incubated at room temperature for 30 min, quenched with 20 mM EDTA and LC-MS confirmed the presence of 2 x 80Da adducts suggesting the ERK2 protein was fully phosphorylated at two sites. The reaction was diluted to ∼400 mL in MonoQ Buffer A and purified as described above; fractions containing ERK2 were pooled, buffer exchanged into SEC buffer and concentrated to 9.6 mg/mL.

#### Phosphorylation of recombinant protein

A solution of 1 mg/mL of recombinant protein was supplemented with 5 mM MgCl_2_, 1 mM ATP (Sigma, A2383) and either 90 nM of recombinant human PKA (PRKACA) kinase (Thermo Scientific, P2912) or 60nM of recombinant ERK2; reactions were incubated at room temperature overnight and phosphorylation states were assessed by LC-MS.

#### Mass spectrometry analysis of tau standards Isoform Quantitation

Total Tau concentrations and relative isoform abundances were determined by targeted liquid chromatography tandem mass spectrometry (LCMS) in a workflow similar to the “FLEXITau” assay described by Mair et al. ^2^. As in their assay, full length stable isotope labeled (SIL) recombinant Tau 2N4R (SILu™ Prot; Sigma; cat #MSST0031) was spiked into samples before trypsin digestion, along with SIL synthetic peptides corresponding to tryptic peptides specific for the 0N, 1N, 2N, 3R, and 4R isoforms. Samples were desalted after digestion by C18 solid phase extraction in pipette tip format (Pierce; cat. #84850). Unmodified tryptic peptides spanning the sequence of Tau plus the isoform-specific peptides were quantified by scheduled, targeted, capillary LC nano-electrospray ionization tandem mass spectrometry in a Thermo Scientific Eclipse orbitrap mass spectrometer. In contrast to the originally reported FLEXITau assay, this acquisition was performed using Parallel Reaction Monitoring (PRM) and ion trap resonance-excitation Collision-Induced Dissociation (CID) rather than Triple Quadrupole Selected Reaction Monitoring with collision-cell CID. Peak areas were extracted from the resulting data using Skyline software ^3^ and normalized to the stable isotope labeled internal standards.

#### Phosphorylation Site Occupancies

Targeted, label-free, middle-down mass spectrometry was performed to estimate phosphorylation site occupancies at T181, S214, S396, S404, and neighboring sites. Samples were prepared as for isoform quantitation, except that before digestion, unmodified lysines were acetylated using acetic anhydride according to previously published methods for histone lysine propionylation^4^. Trypsin digestion thus produced larger peptides with Arg-C specificity, ensuring that the peptides containing T181 and S396 no longer have those phosphorylation sites adjacent to a tryptic cleavage site. Targeted PRM for the masses of each of peptide in its unmodified, singly, doubly, and triply phosphorylated states was performed just as described for isoform quantitation. Peak areas for precursor ions and phosphorylation site-localizing product ions were extracted using the Skyline application. Peak areas at the MS1 level were apportioned to modification status at each phosphorylation site according to the site-specific product ions at the MS2 level, and overall site occupancies calculated with respect to the peak areas covering each site in all of its detected forms.

#### Tau analysis by Western Blot

150 ng of recombinant Tau proteins were spiked into 5 ug Expi293™ extracts, and separated by 4-12% Bolt Bis-Tris Mini Gels (Invitrogen NW04120BOX), then transferred to a PVDF using the iBlot2 Blotting System per manufacturer instructions (Invitrogen Cat #IB24001). Nautilus maintains and provides a Nautilus Probe ID for each unique antibody clone. Membranes were blocked with Pierce Fast Blocking Buffer (Thermo Scientific Cat#37575), probed with primary antibodies including anti-0N (NAUTILUS PROBE ID 21269), anti-2N (NAUTILUS PROBE ID 21269), anti-4R (NAUTILUS PROBE ID 21431), Tau13 (NAUTILUS PROBE ID 21520), HT7 (Invitrogen MN1000), Tau7 (Sigma-Aldrich Mab2239), anti-C-terminal tau (ADx216), anti-pT181 (NAUTILUS PROBE ID 21271), anti-pS202-pT205 (NAUTILUS PROBE ID 21871), anti-pT205 (NAUTILUS PROBE ID 22022), anti-pS214 (NAUTILUS PROBE ID 21273), anti-pT217 (NAUTILUS PROBE ID 22024), anti-pT231 (NAUTILUS PROBE ID 21870), and anti-pS396 (NAUTILUS PROBE ID 21279), then fluorescently labeled secondary antibodies and imaged using Bio-Rad ChemiDoc. Western blot was also used to estimate the abundance of total Tau in cell/tissue extracts by loading Expi293 extracts with different amount of recombinant Tau 0N4R and 2N4R spike-in (ranging from 0.01% to 1%) as references for signal comparison.

#### Cell Culture of Expi293F

Expi293F™ cells were used as a control background for testing of tau-pull-down enrichment as shown in Supplemental Figure 3. Expi293F™ cells (Gibco A14527), grown in suspension as recommended by vendor, were harvest by centrifugation, washed with ice-cold PBS, and lysed by approximately 200 µl RIPA Lysis and Extraction Buffer (Fisher Scientific 89901) supplemented with 1x Protease/Phosphatase Inhibitor cocktail (Cell Signaling Technology #5872) per million cells. Supernatant was collected after 5 min centrifugation at 7,000xg at 4°C, quantified using the Pierce 660nm Protein Assay Reagent following manufacturer’s manual (Thermo Scientific, 22660), and stored at −80C.

#### Cell Culture and Differentiation of iNGN2 Neurons

Details of cell culture reagents, including vendor information and catalog numbers, are described in Shan ^5 6^. Induced neurons (iNGN2) were generated and cultured following established protocols, with brief outlines of cell maintenance, differentiation, and maturation below.

Induced pluripotent stem cells (iPSCs; iP11N, Alstem) were maintained in mTeSR Plus medium (STEMCELL Technologies) supplemented with RevitaCell (1:100; Thermo Fisher), Y-27632 (10 μM; Selleck Chemicals), and doxycycline (3 μg/mL; Sigma-Aldrich). At ∼75% confluency, iPSCs were washed with calcium- and magnesium-free DPBS and incubated with Accutase (STEMCELL Technologies) at 37°C for 10 minutes. Cells were gently dissociated by pipetting, filtered through 40 μm strainers, pelleted at 300 x g for 5 minutes, resuspended in induction medium, counted, and plated at 400,000 cells/cm² on iMatrix-511 (Nippi)-coated vessels (1:200 dilution in DPBS, incubated >1 hour at 37°C).

The induction medium consisted of DMEM/F12 without glutamine (Gibco) supplemented with GlutaMAX (1x), non-essential amino acids (NEAA; 1x), B27 supplement without vitamin A (1x), N2 Max supplement (1x), DAPT (10 μM; Tocris), SB431542 (10 μM; Tocris), Noggin (100 ng/mL; PeproTech), XAV939 (2 μM; Tocris), and doxycycline (3 μg/mL). Medium was refreshed daily from days 1 to 5. On day 5, cytosine β-D-arabinofuranoside (AraC; 5 μM; Sigma-Aldrich) was added to inhibit proliferating cells.

On day 8, neurons were detached with Papain (Worthington Biochemical), dissociated, filtered, and counted. Cells were either cryopreserved in CryoStor CS10 (STEMCELL Technologies) or replated at 200,000 cells/cm² on poly-D-lysine (50 μg/mL; Sigma-Aldrich) and iMatrix-coated vessels for maturation. The maturation medium comprised Neurobasal medium without glutamine (Gibco) supplemented with GlutaMAX (1x), NEAA (1x), B27 with vitamin A (1x), human recombinant BDNF (20 ng/mL; PeproTech), GDNF (20 ng/mL; PeproTech), ascorbic acid (200 μM; Sigma-Aldrich), and dcAMP (500 μM; Sigma-Aldrich). Partial medium changes were performed every 3–4 days.

#### Cell culture and protein extraction of organoids

Organoids were generated at the NeuraCell core facility (Neural Stem Cell Institute, NY, USA). Organoids generated in 96-slitwell plates followed the methods described previously^7^. In brief, when hPSC cultures reached about 70-80% confluency, iPSCs were single cell harvested using TrypLE Express. Cells were counted manually with a hemocytometer and resuspended at approximately 1 million cells/mL in mTeSR1 with 10 µM ROCK inhibitor Y-27632 (Tocris). Each iPSC line was plated into its own 96-slitwell plate (S-bio, MS9096SZ) at a density of 10,000 cells per well to achieve day 1 hPSC spheroids 350-500 µm in diameter. The next day the plate was washed twice, and the medium was then replaced with 14 mL differentiation Medium A: E6 base medium and supplemented with 2.5 µM dorsomorphin (DM), 10 - 20 µM SB431542 (cell-line dependent) and 2.5 µM XAV939. On the sixth day, the medium was changed to Medium B: neural medium (NM) base with soluble growth factors 20 ng/mL EGF plus 20 ng/mL FGF2. Medium was changed daily for 10 days followed by 9 days of feeding three times per week. On day 25, the medium was replaced Medium C comprised of NM supplemented with soluble 20 ng/mL BDNF and 20 ng/mL NT3 and media changes three times per week. From day 43 onward, the medium was changed three times per week with Medium D (NM base with no soluble growth factors) with 15-20 mL per dish (approximately 75% medium changes.) Successful forebrain patterning was achieved for all lines determined by qPCR progenitor expression at 20-day for PAX6 and FOXG1 followed by analysis of 2-month organoid sections by IHC. Organoids were harvested, washed twice with PBS-/-, snapped frozen with liquid nitrogen. To prepare extractions, 3∼4 Organoids were lysed with 75∼100 ul of an in-house RIPA buffer (25mM Tris-HCL pH7.4, 150mM NaCl, 0.1% SDS, 0.5% Na Deoxycholate, 1% IGEPAL CA-630), supplemented with 1x Protease/Phosphatase Inhibitor cocktail (Cell Signaling Technology #5872), clarified by centrifugation at 30,000 g at 4C for 10 minutes, quantified using the Pierce 660nm Protein Assay Reagent following manufacturer’s manual (Thermo Scientific, 22660), and stored at −80C.

### Generation of MiBrains

#### Human iPSC cultures

iPSCs were seeded in Geltrex^TM^ Matrix pre-coated plates and grown as colonies in StemFlex^TM^ medium until they reached 60-70% confluency. At this point, iPSCs were either passaged for maintenance using 0.5 mM EDTA to gently lift colonies, or harvested using Accutase^TM^ cell detachment solution for 5-10 minutes at 37°C to start a differentiation protocol as singularized cells.

#### Differentiation of human iPSCs into neurons

Neuron differentiation was adapted from Zhang et al.^8^ and Lam et al. ^9^. Briefly, iPSCs were transfected with PiggyBac plasmids to confer doxycycline-inducible expression of the Neurogenin-2 gene (*NGN2*, Addgene Plasmid #209077) alone or combination with *SNCA*-A53T-sfGFP (Addgene Plasmid: 209080) or A53T-NAC-SNCA-sfGFP (Addgene Plasmid: 209081), using Lipofectamine^TM^ Stem Transfection Reagent. Briefly, dissociated iPSCs were plated at ∼104,000 cells/cm2 onto Geltrex^TM^-coated plates, in StemFlex™ supplemented with 10 µM Y27632 and 5 µg/mL doxycycline (day 0). At day 1, medium was replaced with Neurobasal N2B27 medium (Neurobasal, 1x B-27, 1x N-2, 1x MEM-NEAA, 1x GlutaMAX, 1% penicillin-streptomycin) supplemented with 10µM SB431542, 100 nM LDN, 5 µg/mL doxycycline, 5 µg/mL blasticidin. At day 2, medium was replaced with Neurobasal N2B27 media supplemented with 10uM SB431542, 100 nM LDN, 5 µg/mL doxycycline, 1 µg/mL puromycin. At days 3-6, medium was replaced daily with Neurobasal N2B27 media supplemented with 5 µg/mL doxycycline, 1 µg/mL puromycin. At day 7, cells were dissociated with accutase and seeded on Poly-L-Ornithine and Laminin pre-coated plates at 156,250 cells/cm2, in Neurobasal N2B27 with 5 µg/mL doxycycline and 10 µM Y27632. At day 8, wells were gently topped with Neurobasal N2B27 supplemented 20ng/mL BDNF, 20ng/mL GDNF, 1mM dcAMP, 2ug/mL Laminin, 1uM AraC, using the same volume of medium in the wells. At day 11, media was replaced with Neurobasal N2B27 supplemented with 10ng/mL BDNF, 10ng/mL GDNF, 0.5 mM dcAMP, 1ug/mL Laminin. This medium was used for half media changes every 3-4 days.

#### Differentiation of human iPSCs into astrocytes

Astrocytes were generated using previously published protocols for iPSC-derived NPC^10^ and astrocyte ^11^ differentiation. Briefly, dissociated iPSCs were plated at 100,000 cells/cm2 onto Geltrex^TM^-coated plates, in pre-warmed StemFlex™ supplemented with 10 µM Y27632. Cells were fed every other day with StemFlex™ until they reached >95% confluence. Once cells reached confluence, the medium was replaced with NPC medium 1:1 DMEM/F12: Neurobasal Medium, 1x N-2 Supplement, 1x B-27 Serum-Free supplement, 1x GlutaMAX Supplement, 1x MEM-NEAA, 1% penicillin-streptomycin) supplemented with 10 µM SB43152 and 100 nM LDN193189 (day 0). From days 1 to 9, cells are fed daily with NPC medium plus 10 µM SB43152 and 100 nM LDN193189. At day 10, cells were split with accutase and replated onto fresh Geltrex^TM^-coated plates, in NPC media supplemented with 20 ng/mL bFGF and 10 µM Y27632. From days 11 to 13, cells were fed with NPC media plus 20 ng/mL bFGF. At day 14, cells were split with accutase and re-seed onto fresh Geltrex^TM^-coated plates, in NPC media plus 20 ng/mL bFGF and 10 µM Y27632. Starting from day 15, cells were fed every 2-3 days with Astrocyte Medium (AM, ScienCell) and passaged using accutase once they reached 90% confluence. From this point, NPCs were fully differentiated into astrocytes in 30 days. NPCs and fully differentiated astrocytes were cryopreserved in freezing medium consisting of 90% knockout serum replacement (KSR) and 10% dimethyl sulfoxide (DMSO).

#### Differentiation of human iPSC into brain microvascular endothelial cells

Brain endothelial cell differentiation was adapted from the protocols from Blanchard et al. ^12^, Qian et al. ^13^ and Wang et al. ^14^. Briefly, iPSCs were transfected with a PiggyBac plasmid to confer doxycycline-inducible expression of the ETS variant transcription factor 2 (ETV2, Addgene Plasmid #168805), using Lipofectamine^TM^ Stem Transfection Reagent. Inducible ETV2-iPSCs were grown until 60-70% confluency, dissociated with Accutase and plated at 20,800 cells/cm2 onto Geltrex^TM^-coated plates in StemFlex™ supplemented with 10 µM Y27632 (day 0). At day 1, medium was replaced with DeSR1 medium (DMEM/F12 with GlutaMAX, 1× MEM-NEAA, 1× penicillin-streptomycin) supplemented with 10 ng/mL BMP4, 6 µM CHIR99021, and 5 µg/mL doxycycline. At day 3, medium was replaced with DeSR2 medium (DeSR1 media, 1x N-2, 1× B-27) supplemented with 5 µg/mL doxycycline. At days 5 and 7, medium was replaced with hECSR medium (Human Endothelial Serum-free Medium, Gibco, 1× MEM-NEAA, 1× B-27, 1% penicillin-streptomycin) supplemented with 50 ng/mL VEGF-A, 2 μM Forskolin, and 5 µg/mL doxycycline. At day 8, cells were dissociated using Accutase and re-seed onto fresh Geltrex^TM^-coated plates in hECSR supplemented with 50 ng/mL VEGF-A and 5 µg/mL doxycycline. This medium was used for every 2-3 days medium change to maintain cells for up to 1 week until ready for tissue assembly (miBrain or JANs).

#### Differentiation of human iPSCs into pericytes

Pericytes were differentiated using previously published protocol from Patsch et al. ^15^. Dissociated iPSCs were plated at 37,000 to 40,000 cells/cm^2^ onto GeltrexTM-coated plates, in StemFlex™ supplemented with 10 µM Y27632 (day 0). At day 1, medium was replaced with N2B27 medium (1:1 DMEM/F12: Neurobasal media, 1x B-27, 1x N-2, 1x MEM-NEAA, 1x GlutaMAX, 1% penicillin-streptomycin) supplemented with 25 ng/mL BMP4 and 8 μM CHIR99021. At days 3 and 4, medium was replaced with N2B27 media supplemented with 10 ng/mL Activin A and 10 ng/mL PDGF-BB. At day 5, pericytes were dissociated with accutase and re-seeded onto fresh 0.1% gelatin-coated plates at 35,000 cells/cm2, in N2B27 supplemented with 10 ng/mL PDGF-BB. This medium was used for every 2-3 days medium change for another 5–7 days. Cells were then banked in freezing medium (90% KSR/ 10% DMSO) and expanded in N2B27 until ready for tissue assembly (miBrain or JANs).

#### Differentiation of human iPSCs into oligodendrocyte progenitor cells

OPC differentiation was adapted from Douvaras et. al., ^16^. Briefly, iPS cells were dissociated into single cells using accutase and seeded at near-confluent density. Differentiation began the next day (designated as day 0) by culturing the cells in DMEM/F12 (1:1) medium supplemented with N2, 10 μM SB431542, 100 nM LDN 193189, and 100 nM all-trans retinoic acid (RA), with daily medium changes. On day 8, 1 μM SAG was added to the differentiation medium, maintaining the presence of 10 μM SB431542 and 100 nM LDN 193189. By day 12, adherent cells were detached and transferred to low-attachment plates to form cell spheres. These spheres were cultured in DMEM/F12 (1:1) medium containing N2, RA, and SAG. On day 30, spheres were plated onto poly-L-ornithine/laminin-coated plates to allow cells to migrate outward. At this stage, the medium was replaced with DMEM/F12 (1:1) supplemented with N2, B27, 10 ng/ml PDGF-AA, 10 ng/ml IGF, 5 ng/ml HGF, 10 ng/ml NT3, 25 μg/ml insulin, 100 ng/ml biotin, 1 μM cAMP, and 60 ng/ml T3. By day 75, cells were harvested, dissociated, and purified using NG2-specific magnetic-activated cell sorting (MACS). The enriched cells were expanded in DMEM/F12 (1:1) medium supplemented with N2, B27, 10 ng/ml PDGF-AA, 10 ng/ml β-FGF, and 10 ng/ml NT3. To drive oligodendrocyte maturation, cells were cultured in DMEM/F12 (1:1) medium supplemented with N2, B27, 20 μg/ml ascorbic acid, 10 mM HEPES, 25 μg/ml insulin, 100 ng/ml biotin, 1 μM cAMP, and 100 ng/ml T3 for at least two weeks.

#### Differentiation of human iPSCs into microglia

iPSC-derived microglia were differentiated via an intermediate differentiation step into hematopoietic progenitor cells (HPCs). For the generation of HPCs the STEMdiff Hematopoietic Kit (cat#: 05310; StemCell Technologies) was used, according to the manufacturer’s manual. Briefly, when 70% confluent (day 0), iPSCs were harvested and passaged at a density of 20-40 colonies per well in a 6-well coated with 0.1mg/mL Matrigel (cat#: 354234; Corning). On day 1, Medium A was added to the culture and on day 4 it was switched to Medium B until complete HPC maturation on day 11-13. Fully differentiated HPCs, floating in the medium and detached from the colonies, were collected for microglial differentiation or frozen in Stem-CellBanker (cat#: 11924; AMSBIO) supplemented with microglial cytokines. For the generation of mature microglia, differentiated HPCs were collected and transferred into Matrigel-coated 6-well plate at a confluency of 350k HPCs per well. The differentiation takes 25-28 days, during which HPCs are cultured in microglia media consisting of DMEM/F12 (cat#: 11320-033; Gibco), with 2X B27 (cat#: 17504044; Thermo Fisher Scientific), 0.5X N2 (cat#: 17502048; Thermo Fisher Scientific), 1X Glutamax (cat#: 35050061; Gibco), 1X non-essential amino acids (cat#: 11-140-050; Gibco, 400 mM Monothioglycerol (cat#: M6145; Millipore Sigma), and 5 mg/mL human insulin (cat#: I9278; Millipore Sigma), freshly supplemented with 100 ng/mL IL-34 (cat#: 200-34; PeproTech), 50 ng/mL TGFβ1 (cat#: 100-21; PeproTech), and 25 ng/mL M-CSF (cat#: 300-25; PeproTech). Microglia was added to miBrains within the pool of geltrex-encapsulated cells at the time of miBrain assembly. MiBrains were maintained in miBrain media supplemented with 100 ng/mL IL-34 and 25 ng/mL M-CSF for 1 week, and then switched to miBrain media supplemented with 25 ng/mL M-CSF until downstream experiments.

#### 3D Tissue Assembly for miBrains

Neurons, astrocytes, endothelial cells, pericytes, and OPCs were dissociated using accutase or TryplE Select (astrocytes). Cells were resuspended in corresponding media, counted, and resuspended at 1 x 106 cells/ mL. For miBrains, a tube was prepared to contain 5 x 106 neurons, 5 x 106 endothelial cells, 1 x 106 astrocytes, 1 x 106 OPCs, and 1 x 106 pericytes per 1 mL. For neuron + astrocyte co-cultures, a tube was prepared to contain 5 x 106 neurons and 1 x 106 astrocytes per 1 mL. Pooled cells were spun down at 200 x g for 5 min at RT. Media was aspirated carefully, leaving the cell pellet undisturbed. The cell pellet was placed on ice and resuspended in 1 mL Geltrex^TM^ supplemented with 10 µM Y27632 and 5 µg/mL doxycycline, avoiding air bubbles and keeping it on ice to prevent premature Geltrex polymerization and inability to seed miBrains or co-cultures properly. To generate miBrain tissue that adopted a free-floating, organoid-like morphology over time, 25-50 µL of encapsulated cell suspension were seeded per inner glass-bottom well of a 48-well MatTek plate (MatTek). To generate miBrain tissue that remained attached to the plate (more suitable for automated imaging) while conserving 3D morphology, 10 µL of encapsulated cell suspension were seeded per well of a 96-well µClear plastic-bottom plate (Greiner). For JANs assembly and cryopreservation, a tube containing 5 x 106 endothelial cells, 1 x 106 astrocytes, 1 x 106 OPCs, and 1 x 106 pericytes per 1 mL was prepared. Pooled cells were spun down at 200 x g for 5 min at RT and cryopreserved in miBrain freezing media (60% KSR, 30% hECSR medium, 10% DMSO, 10 µM Y27632, 50 ng/mL VEGF-A). Upon thaw, cell viability was assessed, and the appropriate volume of neurons needed to conserve the original miBrain cell-to-cell ratio was added to the pooled cell suspension, which was spun again, encapsulated, and seeded as described above. After miBrains were seeded, the plates were transferred into a 37 °C 95%/5% Air/CO_2_ incubator for 30 minutes to allow the Geltrex^TM^ to polymerize. After polymerization of the gel, miBrain week-1 medium (Human Endothelial Serum-free Medium, 1x Pen/Strep, 1X MEM-NEAA, 1X CD Lipids, 1x Astrocyte Growth Supplement (ScienCell), 1x B27 Supplement, 10µg/mL Insulin, 1µM cAMP-dibutyl, 50µg/mL Ascorbic acid, 10ng/mL NT3, 10ng/mL IGF, 100ng/mL Biotin, 60 ng/mL T3, 50 ng/mL VEGF, 1µM SAG, 5 µg/mL doxycycline) was added to each well, ensuring complete submersion of the culture in media (250-500uL per well of a 48-well matTek plate, 100 to 200 µL per well of a 96-well plate). Half media change was performed every 2-3 days. On day 8 after miBrain seeding, the media was changed to miBrain week-2 medium (Human Endothelial Serum-free Medium, 1x Pen/Strep, 1X MEM-NEAA, 1X CD Lipids, 1x Astrocyte Growth Supplement (ScienCell), 1x B27 Supplement, 10µg/mL Insulin, 1µM cAMP-dibutyl, 50µg/mL Ascorbic acid, 10ng/mL NT3, 10ng/mL IGF, 100ng/mL Biotin, 60 ng/mL T3, 5 µg/mL doxycycline). Cultures were used for downstream assays after 2 weeks.

#### miBrain protein extraction

*APOE3* and *APOE4* genotype miBrain samples are synthesized from CRISPR-edited iPSC isogenic pairs that differ at the *APOE* locus as previously described ^17,18^. Upon harvest, miBRains were rinsed with PBS then lysed in 400 µL Pierce^TM^ RIPA Buffer (Thermo Scientific, 89900) supplemented with 1X Halt^TM^Protease & Phosphatase Inhibitor Cocktail (Thermo Scientific, 1861281) and 1X 0.5 M EDTA Solution (Thermo Scientific, 1861274). Lysis digestion was facilitated by using a disposable dounce homogenizer and two rounds of trituration with p200 pipettes followed by vortexing. The samples were centrifuged 15,000 rpm for 15 minutes at 4° C. The supernatant was snap frozen with liquid nitrogen and stored at −80C. Protein concentrations were quantified using the Pierce 660nm Protein Assay Reagent following manufacturer’s manual (Thermo Scientific, 22660).

#### Mouse and human brain tissues and protein extraction

Half brains from transgenic mice were obtained from Quest Pharmaceutical Services (Newark, DE). The mice (known as 8c mice) have a hybrid background of C57/blk6, DBA, Swiss Webster, C57BL/6 and 129/SvJae, expressing a human tau derived from a human PAC, H1 haplotype, and with the murine Tau gene knocked out by a targeted disruption of exon1.

A human frontal cortex brain sample (as used in Figure 3) was obtained from Analytical Biological Services Inc. (New Castle, DE). The specimen was collected at 6 hours 50 minutes postmortem. The donor was a Caucasian male at the age of 66 with lung cancer, without neurological disorder.

Brain protein extraction for both mouse and human reference samples was developed based on Ericsson et al., ^19^ with minor modifications. Specifically, one piece of mouse brain hemisphere sectioned along the midline (250 mg) or human brain specimen (∼300 mg) was first homogenized by grinding in a mortar pre-chilled with liquid nitrogen. 9 volumes of brain extraction buffer (2% SDS and 20 mM HEPES pH 6.8) supplemented with 1x Protease/Phosphatase Inhibitor cocktail (Cell Signaling Technology #5872) was added drop wise, then ground into fine powder with the brain sample. The mixture was constantly held at room temperature until melt, then transferred into 2 ml Eppendorf tubes, at about 1 ml per tube, and incubated at 70°C for 10 minutes with constant shaking at 1400 rpm in a Thermomixer. Supernatant was collected after 10 min centrifugation at 14,000 g at room temperature, quantified using the Pierce 660nm Protein Assay Reagent following manufacturer’s manual (Thermo Scientific 22660), and stored at −80C.

The cohort of seven post-mortem human patient brain samples referenced in the manuscript and figures as ‘aged human brains’ in Figure 5 is detailed in **Supplemental Table S6**. Samples were obtained from Mount Sinai Neuropathology Brain Bank and Research CoRE. After thawing from −80C storage, the samples were lysed in 200 µL Pierce^TM^ RIPA Buffer (Thermo Scientific, 89900) supplemented with 1X Halt^TM^Protease & Phosphatase Inhibitor Cocktail (Thermo Scientific, 1861281). Lysis digestion was facilitated by using a disposable dounce homogenizer followed by one round of trituration with a p1000 pipette followed by vortexing for 15 seconds and then another round of trituration with a p200 pipette followed by vortexing for 15 seconds. The samples were centrifuged 15,000 rpm for 15 minutes at 4° C. The supernatant was snap frozen with liquid nitrogen and stored at −80°C. Protein concentrations were quantified using the Pierce 660 nm Protein Assay Reagent following manufacturer’s manual (Thermo Scientific, 22660).

Protein was extracted from these samples with Pierce RIPA buffer (Thermo Scientific 89900), they were first diluted with additional RIPA buffer to 4 mg/ml total protein, then diluted with equal volume of brain extraction buffer (2% SDS and 20 mM HEPES pH 6.8) supplemented with 1x Protease/Phosphatase Inhibitor cocktail (Cell Signaling Technology #5872) to 2mg/ml total protein, then treated with 3 mM of Dithiobutylamine (Sigma Aldrich 774405) for 10 min at 70°C, followed by iodoacetamine (Pierce A39271) at 10 mM for 15 min at 25°C. The protein was precipitated by four volumes of pre-chilled (−20°C) acetone. The pellets were briefly washed with cold acetone, air dried, then resuspended in PBS supplemented with 1% SDS. Lysates were quantified with Pierce 660 Assay, adjusted to 2 mg/ml, then treated with 200 µM d (mTz) with 2% DMSO at 25°C for 3 hours on a thermomixer shaking at 850 rpm protected from light. The reaction was quenched with L-Arginine (Sigma #1100) at 50mM for 30 minutes at 25°C. Modified lysates were diluted by 5 volumes of IP Buffer and processed as described in Methods.

#### Flow cell preparation and assembly

Silicon wafers with nano-scale features patterned in JSR resist were obtained from an outside vendor (Skywater Technologies). The wafers were then functionalized with 3-Aminopropyltrimethoxysilane (APTMS) (Gelest P/N:SIA0611.1) using an IST RPX-540 chemical vapor deposition system (Integrated Surface Technologies). At least one day after APTMS modification, the wafers were immersed for two hours in a 10 mM solution of NHS ester PEG (5k) azide (Jenkem P/N: A5088) in 50% dimethylsulfoxide (Sigma Aldrich, P/N 472301) and 50% water at room temperature. The wafers were rinsed and then immersed in a 0.1 µM solution of DBCO-oligo (Integrated DNA Technologies, /5DBCOTEG/TAAATGCGGGCTAATAACTGT) in 50% dimethylsulfoxide (Sigma Aldrich, P/N 472301) and 50% water at 35°C for two hours. The wafers were then rinsed and dried. The following day, the wafers were coated with Plaincoat (Nikka Seiko Co, P/N EG3-1532) at 500 rpm for 40 seconds on a APOGEE spin coater (Cost Effective Equipment). The wafers were then packaged and shipped for dicing into 40 x 33 mm die (GDSI, San Jose).

Once returned from dicing, the individual dies were removed with tweezers from the dicing tape. They were placed in a stainless-steel rack for resist removal. Resist was removed from the individual dies using a custom wetbench (Amerimade). The rack containing the dies was first soaked in a solution of 1% triethylamine (Fisher Scientific, P/N O4884500) in Remover PG (Kayaku Advanced Materials, P/N G050200) at 50°C for five minutes. The rack of dies was then sonicated for five minutes at 50°C in the same bath. The rack was then moved to a fresh tank of 1% triethylamine in Remover PG and sonicated for an additional five minutes at 50°C. The rack of dies was then transferred to a third bath where the dies were soaked for one minute in isopropyl alcohol. Finally, the rack was transferred to a nitrogen dryer, where the dies were dried under hot nitrogen for twenty minutes.

The dies were then assembled into flow cells, using adhesive (Adhesives Research P/N: 92712) with 4 defined lanes per flow cell. The adhesive was attached to a 170 micron thick glass backer (40mm x 33 mm) made of D263 (Corning glass, cut to size by Citrogene, Inc). Before assembly, the D263 glass backer was cleaned in a solution of 1% CIP100 (Steris Life Sciences) in DI water for 15 minutes at 70°C, and then washed with DI water for 3 minutes and dried with nitrogen. The backers were then coated with Hexamethyldisilazane (HMDS) via chemical vapor deposition at Integrated Surface Technologies. The assembled flow cell was then glued into a custom-machined aluminum caddy (V-Tech Manufacturing, Inc.). The assembled flow cells were then vacuum sealed into nitrogen purged PAKDRY7500 mylar foil bags (IMPAK).

## Supplemental Tables

**Supplemental Table 1:**
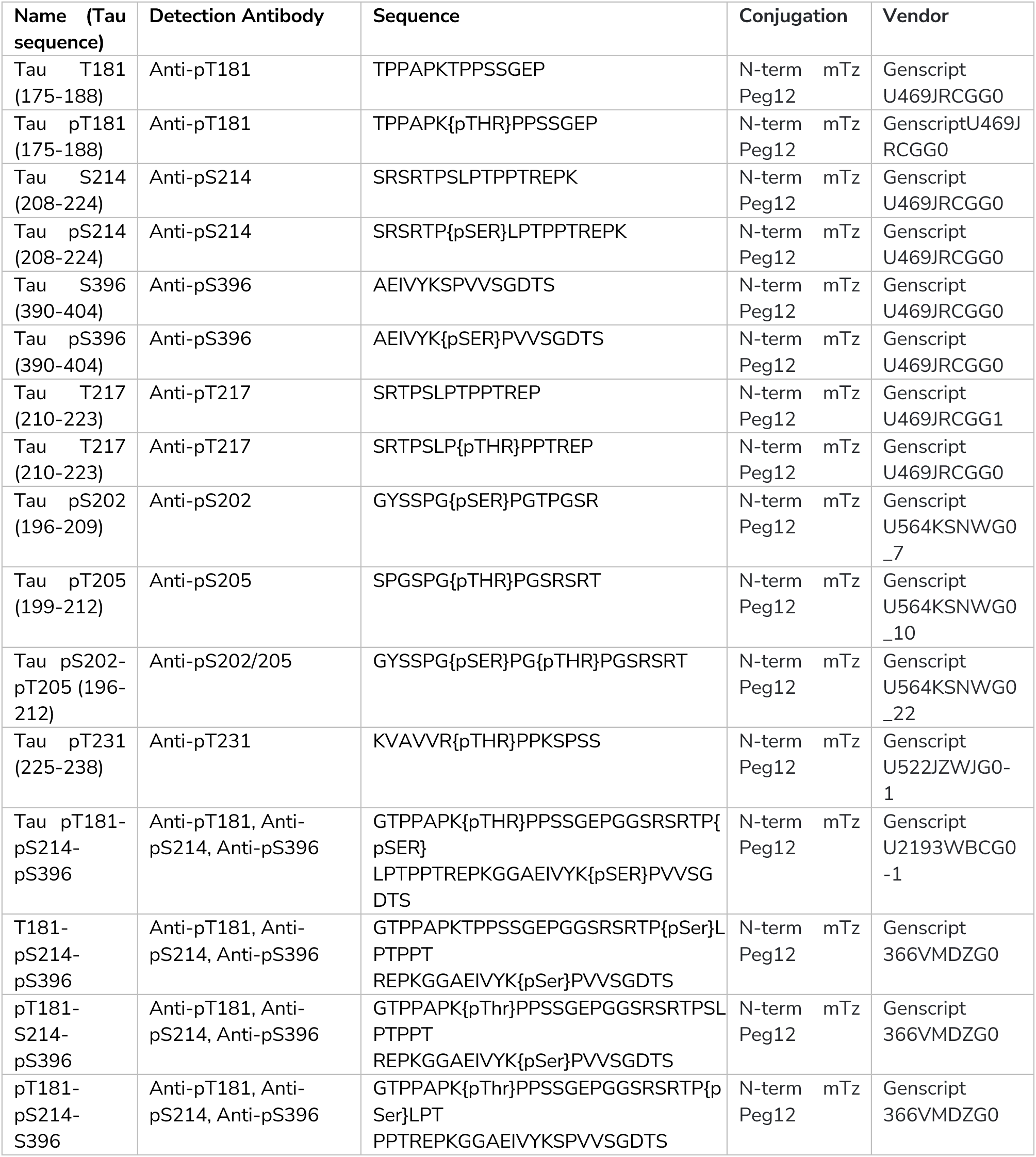
Synthetic, mTZ modified peptides used for antibody suitability screening.

**Supplemental Table S2:**
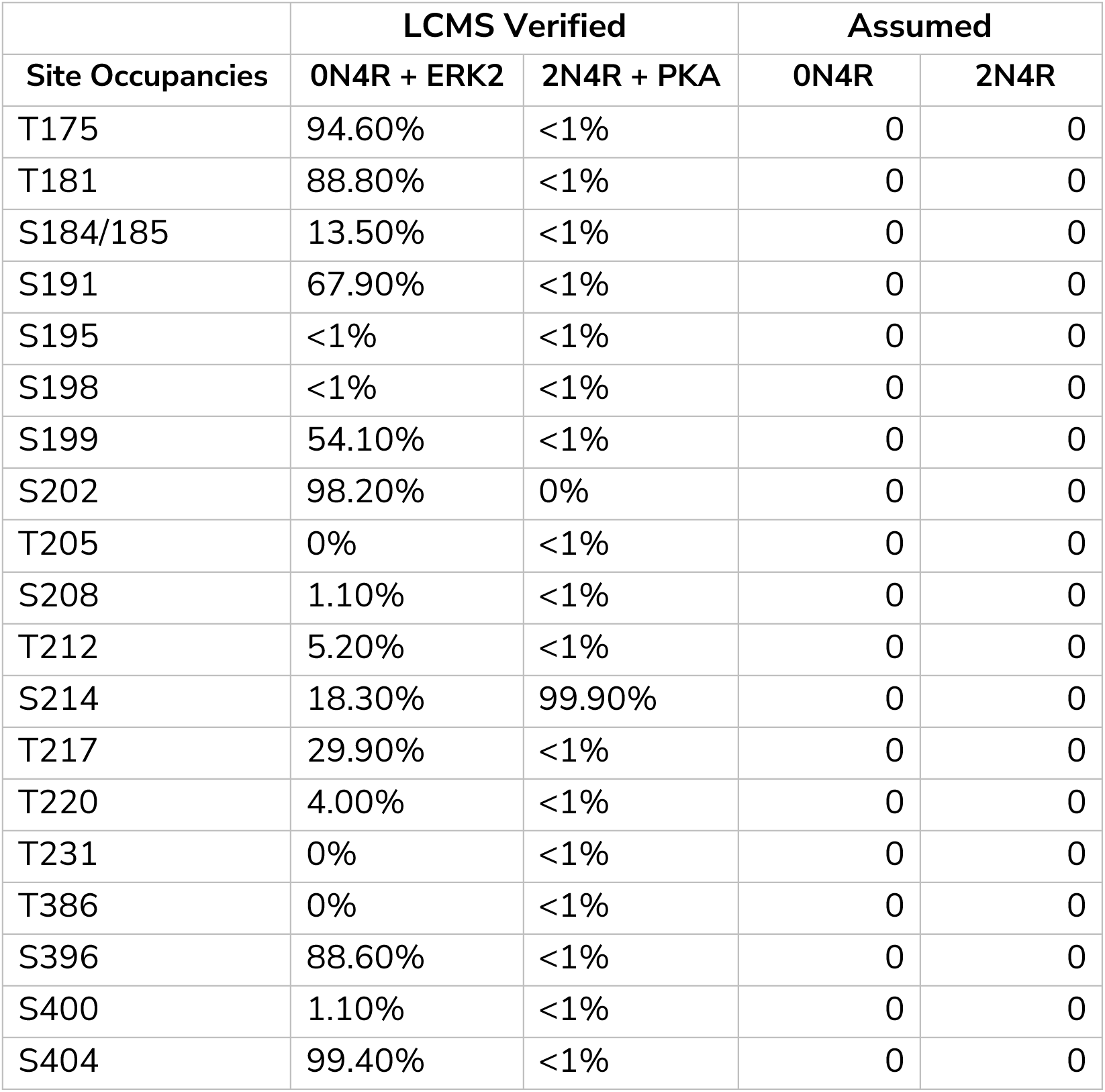
Site occupancy and percent phosphorylation of Tau protein standards as assessed by LCMS. Phosphorylation site occupancy provides the ground truth for the presence of absence of phosphorylation in our protein standard with Tau 0N4R and 2N4R isoforms showing no detectable phosphorylation, while 0N4R modified with ERK2 showed multiple phosphorylation across the backbone at various occupancy and the 2N4R isoform modified with PKA showed almost complete phosphorylation at serine 214.

**Supplemental Table S3:**
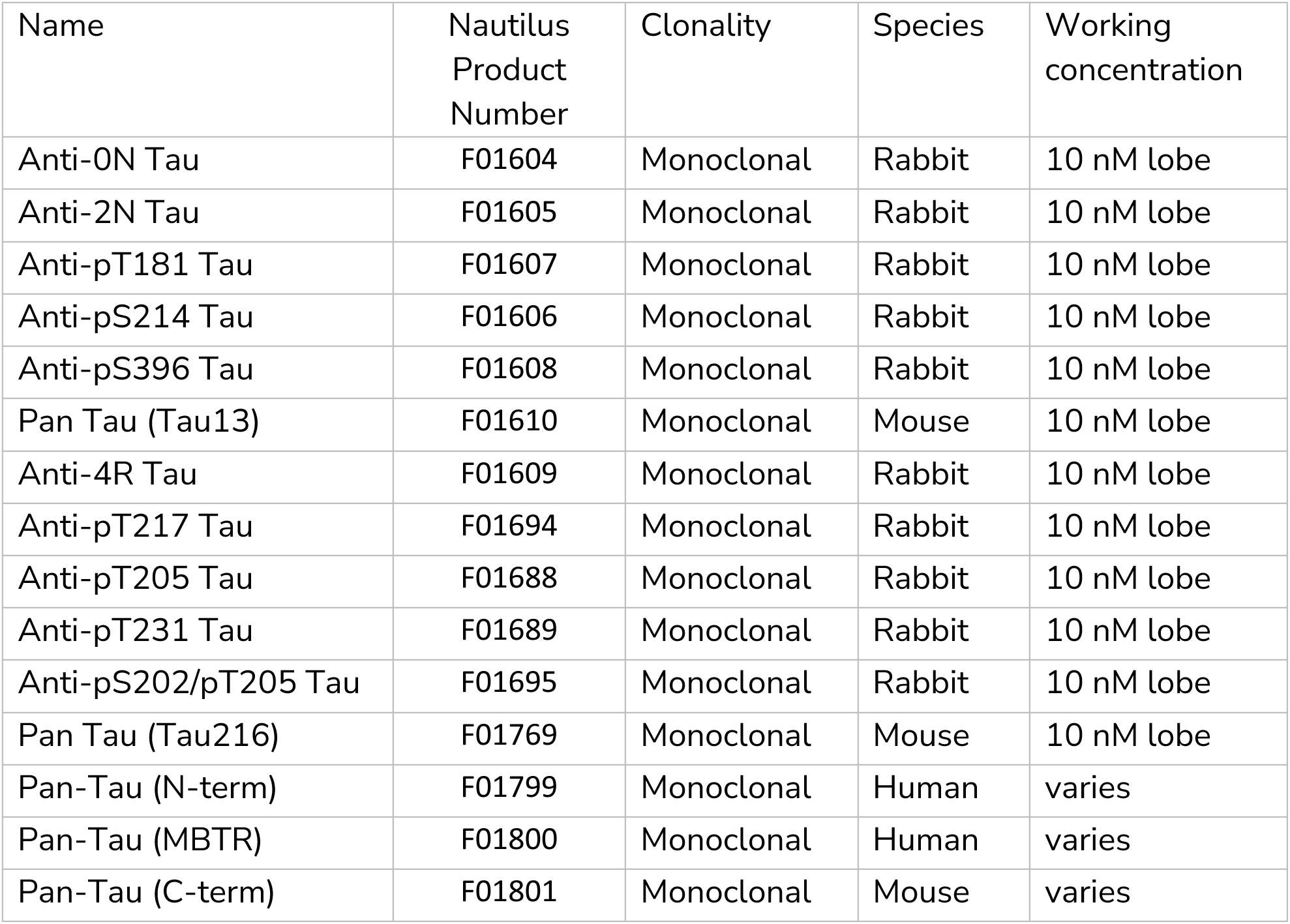
List of labeled-antibody-conjugates used in the single-molecule assay and for Tau enrichment.

| Name | Nautilus Product Number | Clonality | Species | Working concentration |
| --- | --- | --- | --- | --- |
| Anti-0N Tau | F01604 | Monoclonal | Rabbit | 10 nM lobe |
| Anti-2N Tau | F01605 | Monoclonal | Rabbit | 10 nM lobe |
| Anti-pT181 Tau | F01607 | Monoclonal | Rabbit | 10 nM lobe |
| Anti-pS214 Tau | F01606 | Monoclonal | Rabbit | 10 nM lobe |
| Anti-pS396 Tau | F01608 | Monoclonal | Rabbit | 10 nM lobe |
| Pan Tau (Tau13) | F01610 | Monoclonal | Mouse | 10 nM lobe |
| Anti-4R Tau | F01609 | Monoclonal | Rabbit | 10 nM lobe |
| Anti-pT217 Tau | F01694 | Monoclonal | Rabbit | 10 nM lobe |
| Anti-pT205 Tau | F01688 | Monoclonal | Rabbit | 10 nM lobe |
| Anti-pT231 Tau | F01689 | Monoclonal | Rabbit | 10 nM lobe |
| Anti-pS202/pT205 Tau | F01695 | Monoclonal | Rabbit | 10 nM lobe |
| Pan Tau (Tau216) | F01769 | Monoclonal | Mouse | 10 nM lobe |
| Pan-Tau (N-term) | F01799 | Monoclonal | Human | varies |
| Pan-Tau (MBTR) | F01800 | Monoclonal | Human | varies |
| Pan-Tau (C-term) | F01801 | Monoclonal | Mouse | varies |

**Supplement Table S4:**
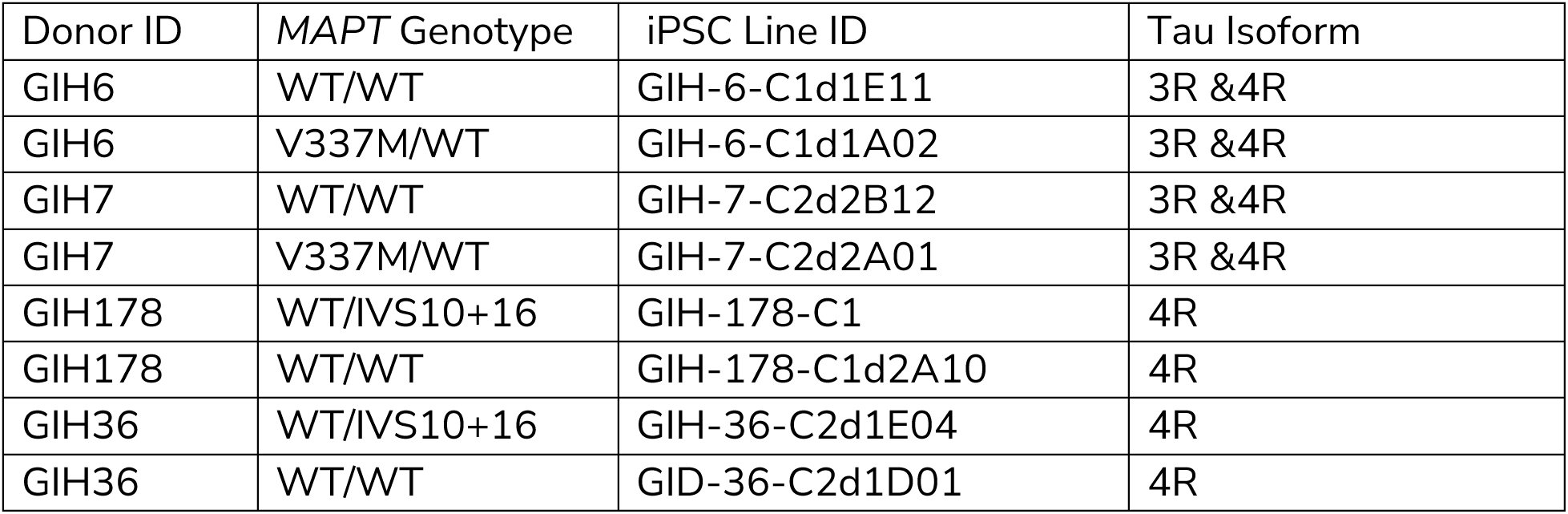
Cortical organoid genotype, donor and iPSC line identification^20^. Patient-derived organoids and gene-edited control cell lines used for Tau proteoform characterization. The table links donor ID, *MAPT* genotype and associated molecular Tau changes.

**Suppl. Table S5:** Number of proteoforms found in 1 or more model systems. Number of proteoforms detected above 0.1% abundance in at least one sample and shared across the five tissue types investigated.

| Number of Model Systems | Not Phosphorylated | Phosphorylated | Total |
| --- | --- | --- | --- |
| 1 | 0 | 34 | 34 |
| 2 | 2 | 39 | 41 |
| 3 | 0 | 23 | 23 |
| 4 | 0 | 21 | 21 |
| 5 | 4 | 7 | 11 |
| Total across all model systems | 6 | 124 | 130 |

**Supplemental Table S6:**
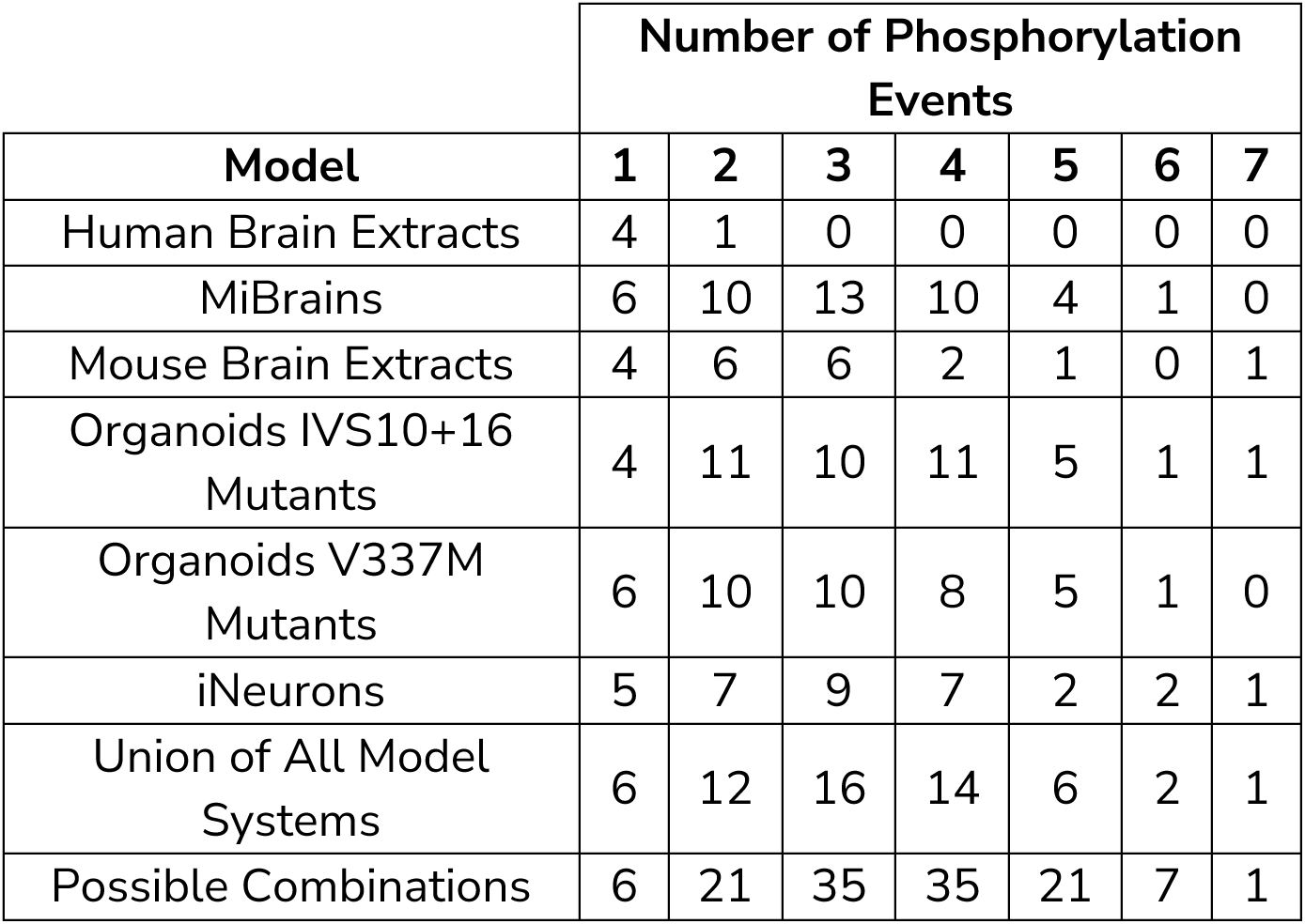
Number of phosphorylation patterns observed and possible.

**Supplemental Table S7:** Characteristics of brain samples used in Figure 5.

| NPBB# | Age at Death | Sex | ABC Score | Lewy Bodies? | APOE Genotype | Notes |
| --- | --- | --- | --- | --- | --- | --- |
| 127 | 70 | M | NA | NA | E3/E3 | Cognitively normal |
| 204 | 65 | F | NA | NA | E3/E3 | Cognitively normal |
| 203 | 92 | F | A2B2C3 | No | E3/E3 | Impaired |
| 122 | 72 | M | A2B3C2 | Yes | E4/E4 | Impaired |
| 213 | 96 | M | A2B3C3 | Yes | E3/E3 | Impaired |
| 18 | 83 | F | A3B3C3 | No | E3/E4 | Impaired |
| 12 | 78 | F | A2B3C2 | No | E3/E4 | Impaired |

## Supplemental Figures

**Suppl. Fig. 1:**
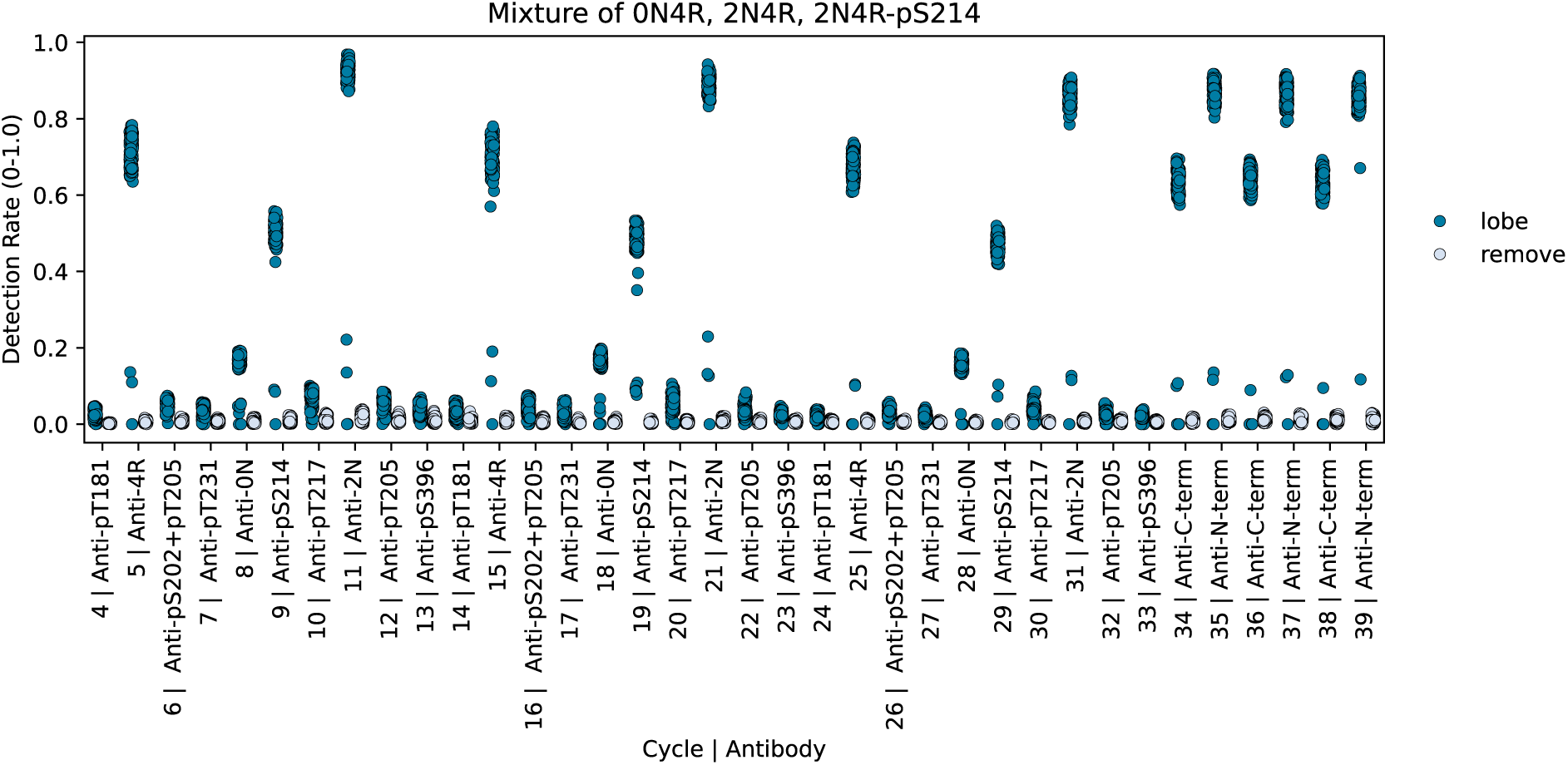
Method of iterative mapping of proteoforms. Representative co-localization plot of antibody binding and removal over 36 cycles using a panel of 12 antibodies. Y-axis shows observed rate of co-localization of fluorescence associated with labeled probes at 647 nm with labeled protein nanoparticle-conjugate at 488 nm through 36 cycles (x-axis). Epitope binding events are indicated in dark blue and light blue dots show fluorescence following removal of antibody nanoparticle conjugates.

**Suppl. Fig. 2:**
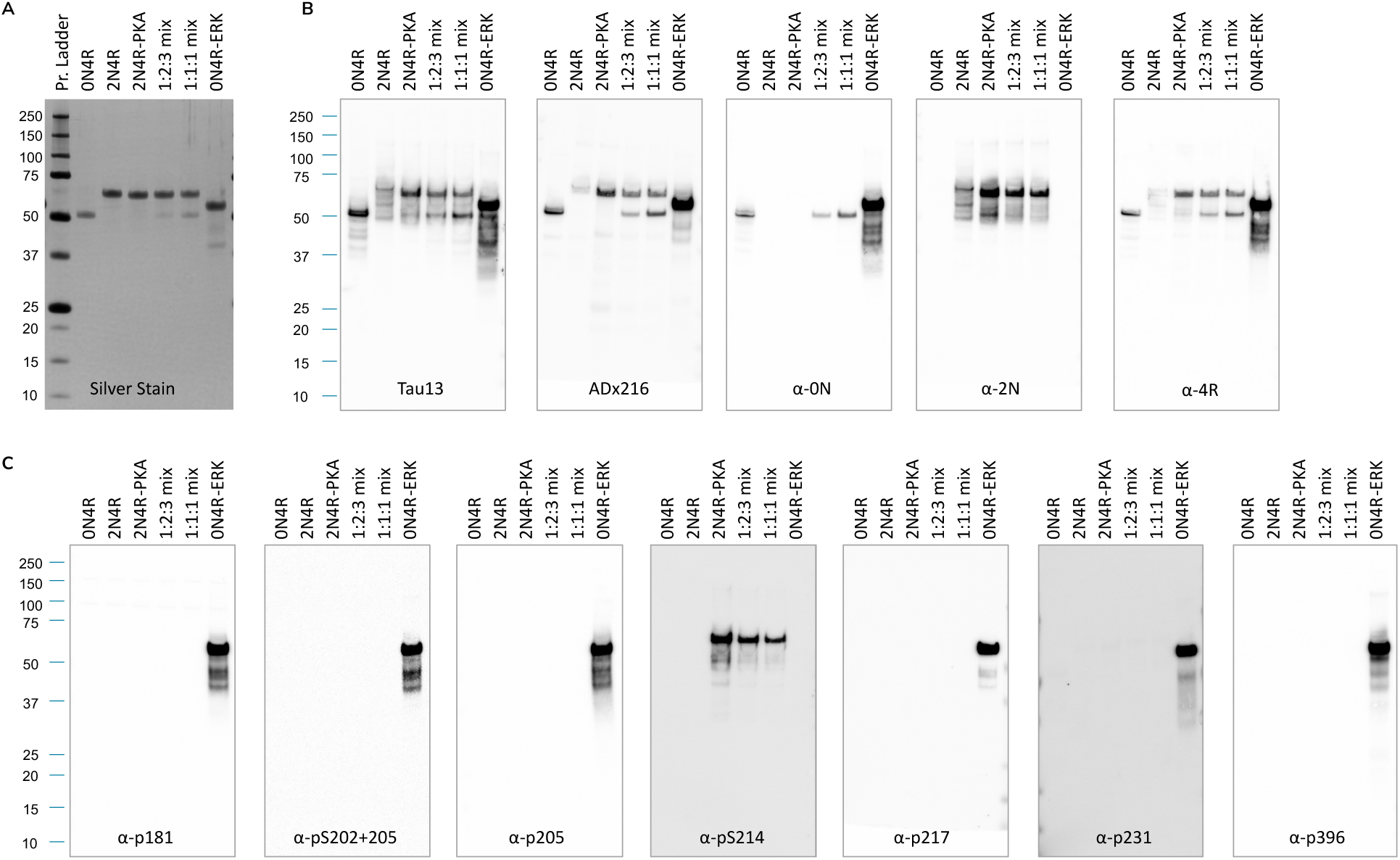
Western blot is unable to resolve proteoform complexity. (A) Analysis of a mixture of Tau proteoforms using silver staining showing individual proteoform standards in lane 2-4 and 7, and a mixtures of three proteoforms in lane 5-6 (0N4R, 2N4R, and 2N4R-PKA mixed at ratios of 1:2:3 and 1:1:1, respectively). (B) Western blots using pan Tau antibodies (Tau13 and Adx216), and isoform specific antibodies (anti-0N, −2N and −4R). (C) Western blot using phosphorylation specific antibodies used in this study.

**Suppl. Fig. 3.**
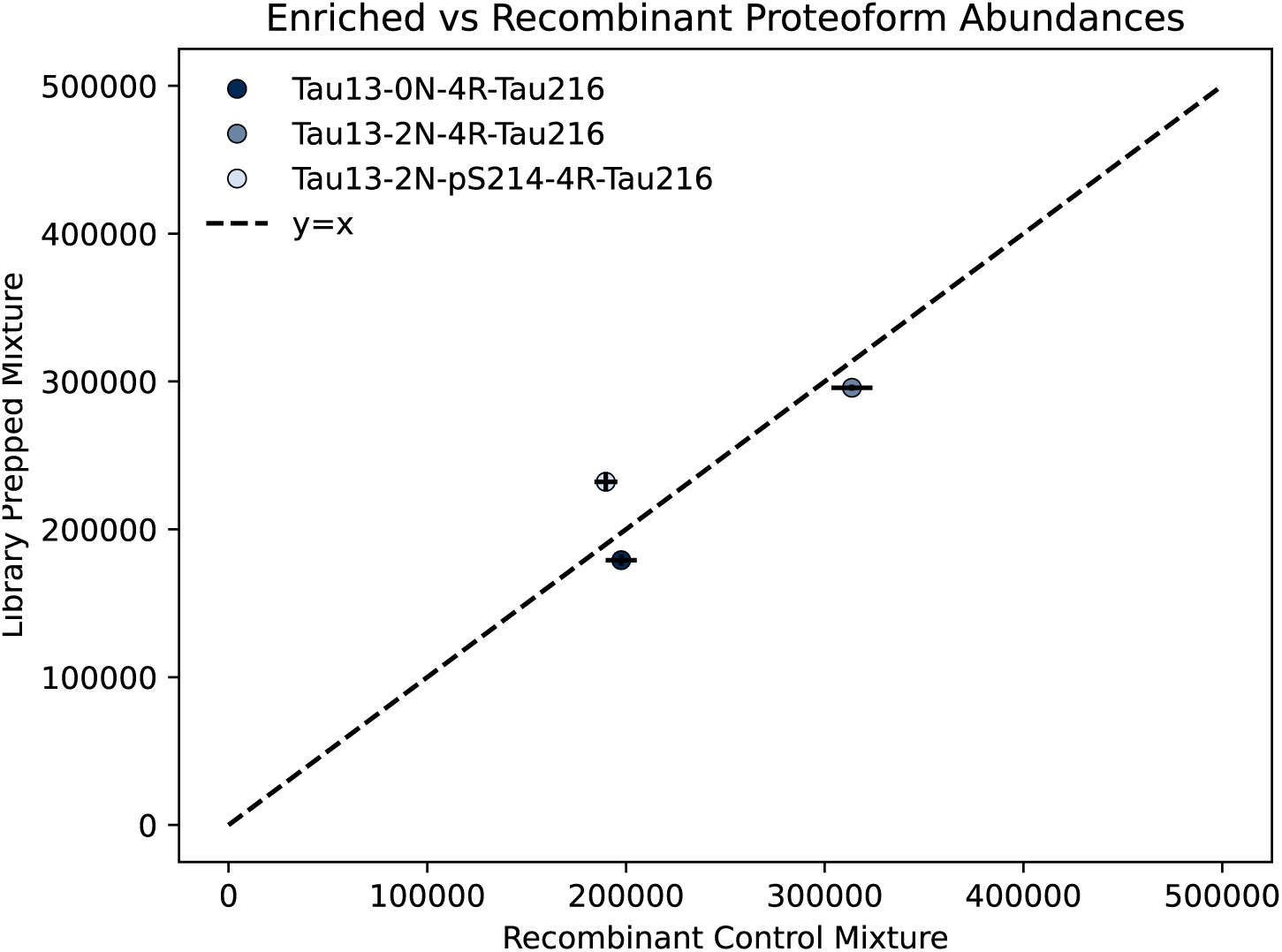
Tau proteoform enrichment bias test comparing measured signal of proteoforms standard mixture against expected. (0N4R, 2N4R, 2N4R-PKA) spiked into a Expi293 cell lysate, under enriched (x-axis) and unenriched (y-axis) samples. Each point represents a distinct tau proteoform, plotted according to its signal intensity in the unenriched (x-axis) versus enriched (y-axis) samples.

**Suppl. Fig. 4A:**
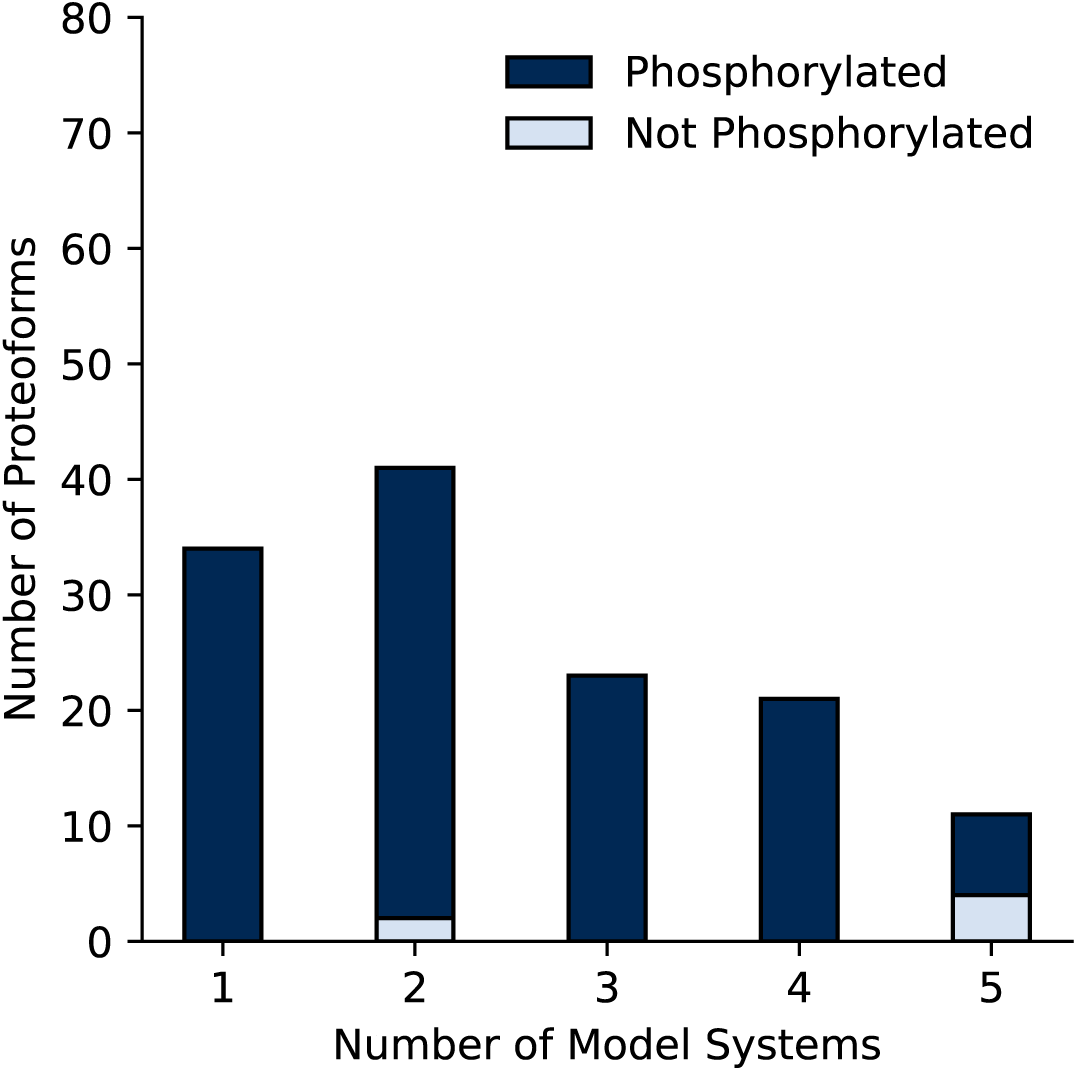
Number of proteoforms found in 1 or more model systems. Number of full length tau proteoforms detected above 0.1% abundance and shared across the five neuronal tissues investigated (see Supplemental table S6).

**Suppl. Fig. 4B:**
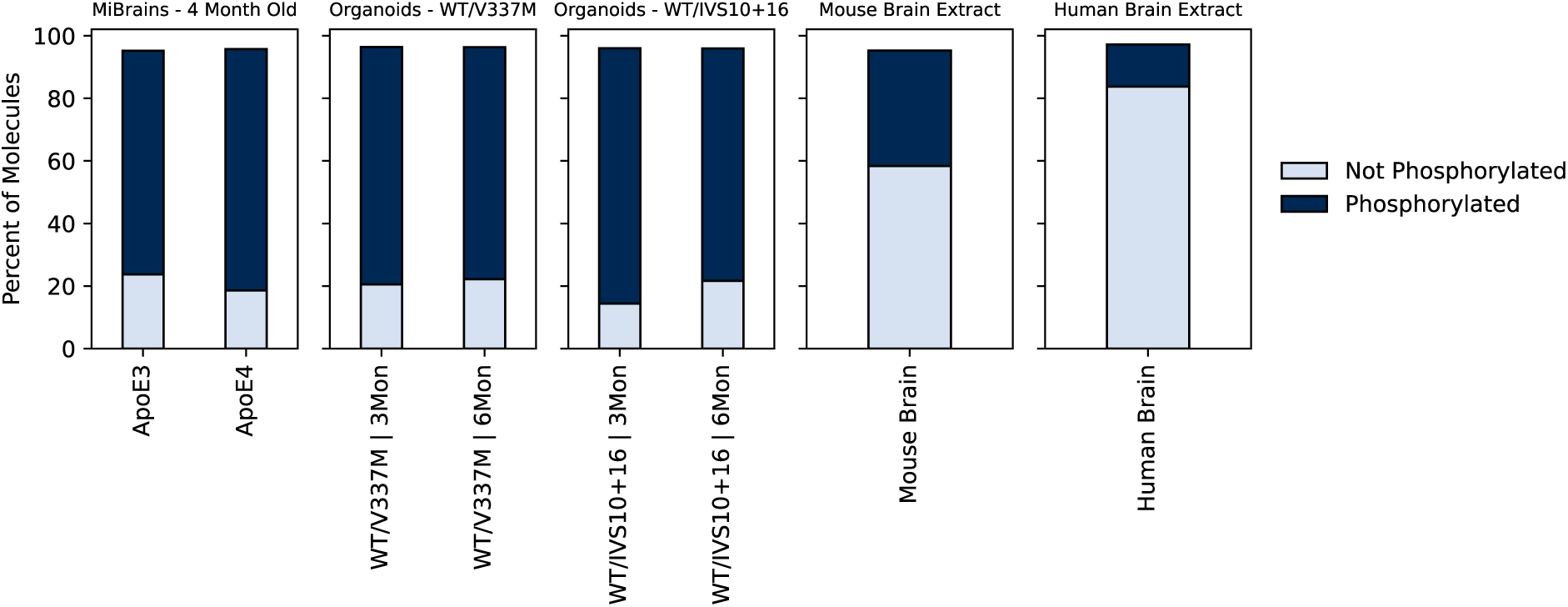
Percent tau protein molecules phosphorylated. The fraction of tau proteoforms modified by one or several phosphorylations for samples extracted from miBrains (APPOE3 genotype), cortical organoids (WT/V337M and WT/ISV10+16 mutants) at 3 and 6 months maturity, hTau mouse brain and human brain extract.

**Suppl. Fig. 4C:**
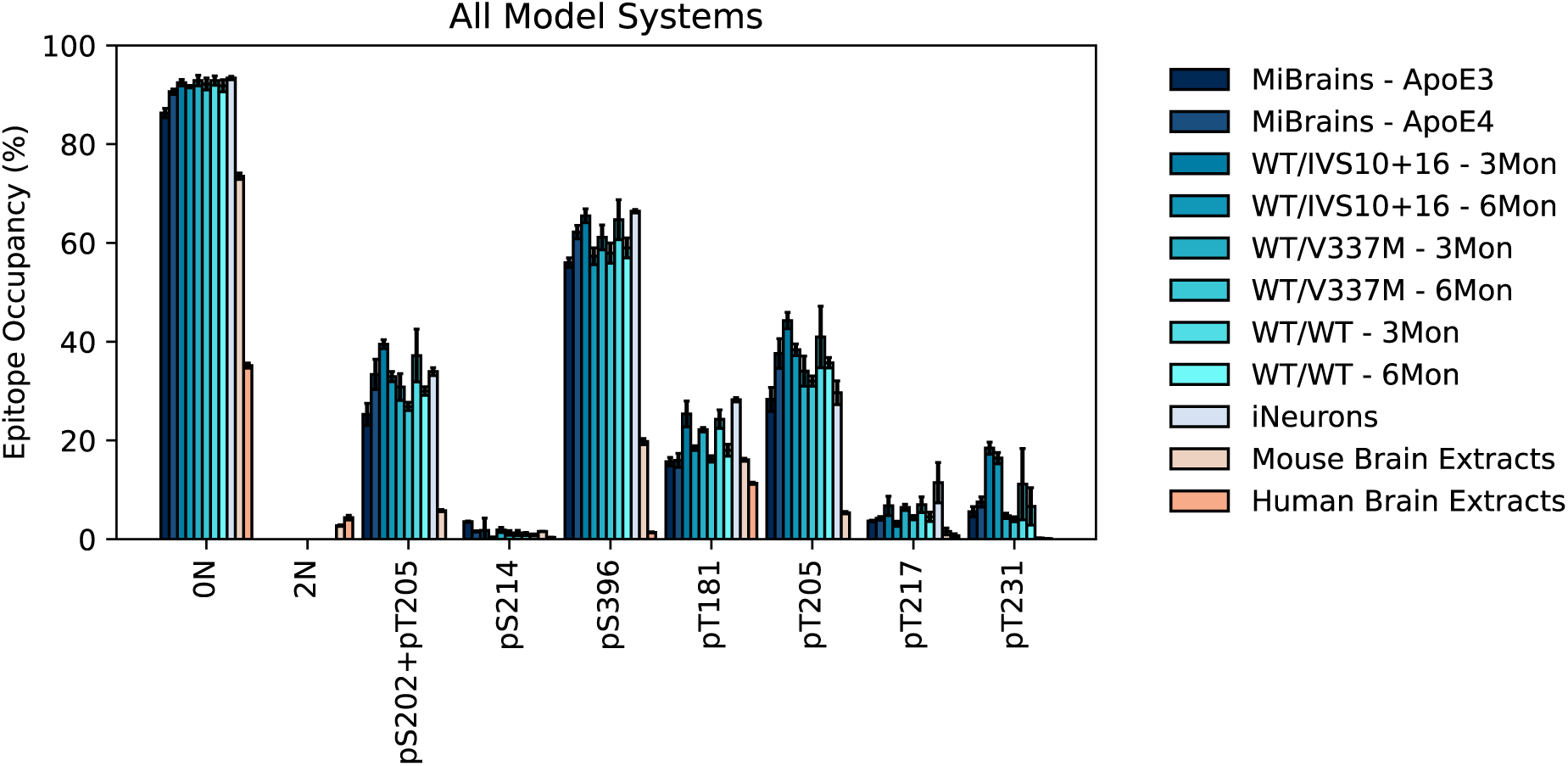
Prevalence of isoforms and occupancy of individual PTM sites found in cellular and animal model systems. Tau proteoforms were extract from human brain extract, mouse brain, miBrains, cortical organoids (WT/V337M and WT/IVS10+16 mutant) and iNeurons.

**Suppl. Fig. 4D:**
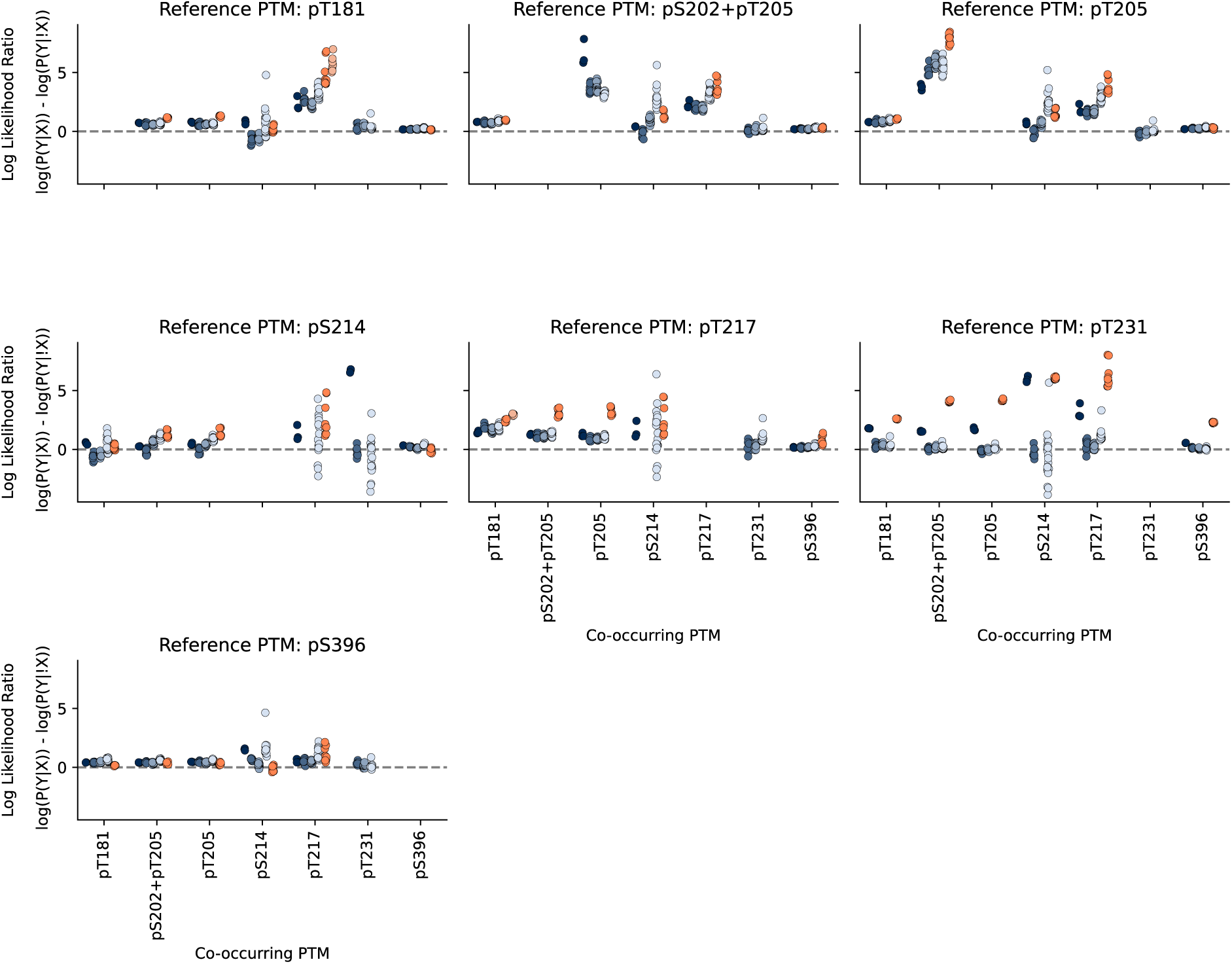
Log likelihood ratio of co-occurrence rate for all proteoforms observed. Rate of co-occurrence at which post translational modification Y occurs given post translational modification X to the rate at which post translational modification Y occurs when post translational modification X is not present. Each data point represents a sample.

**Suppl. Fig. 4E:**
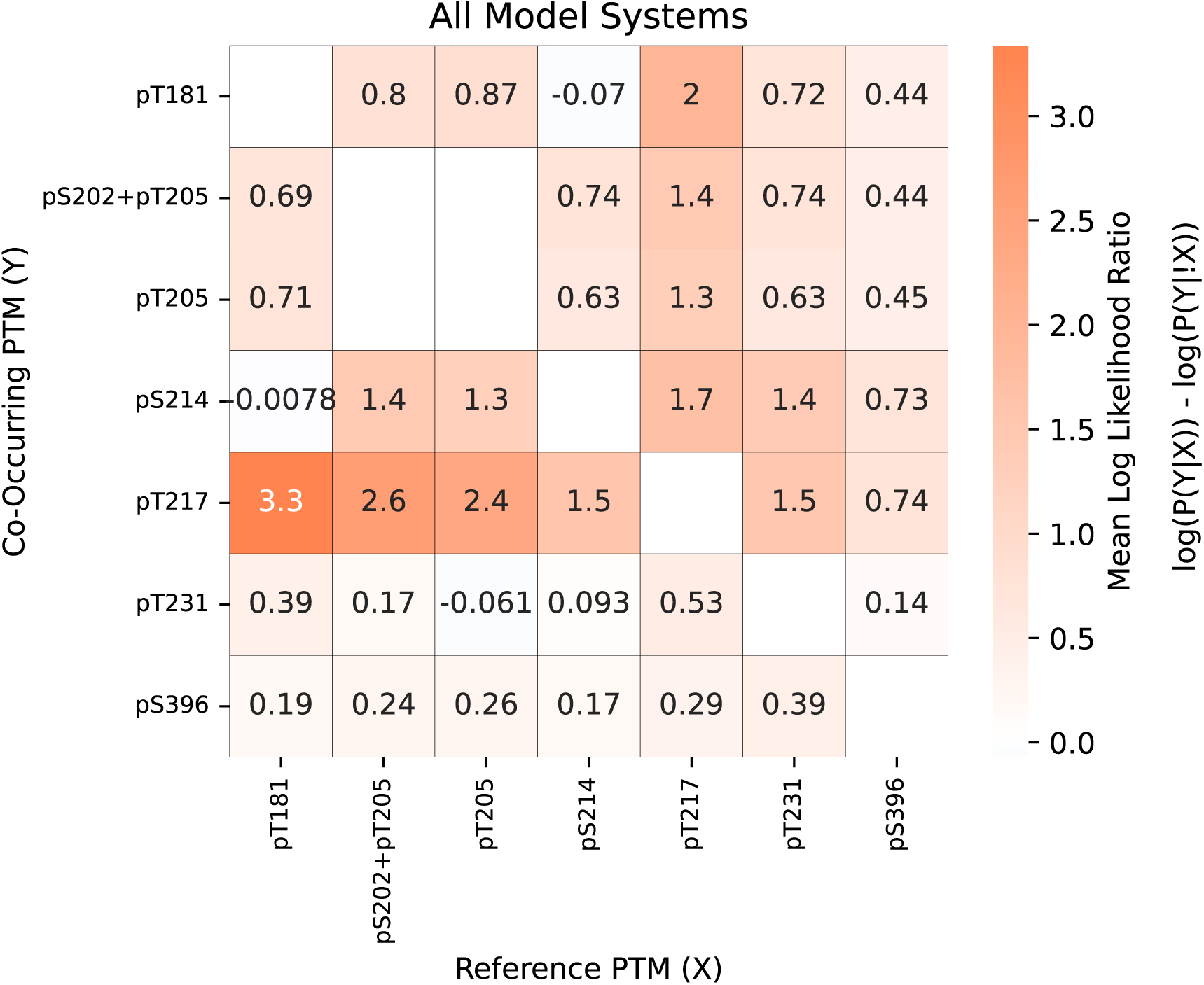
Median Log likelihood ratio of co-occurrence rate. Rate of co-occurrence at which post translational modification Y occurs given post translational modification X to the rate at which post translational modification Y occurs when post translational modification X is not present. Median value across all model systems.

**Suppl. Fig. 5:**
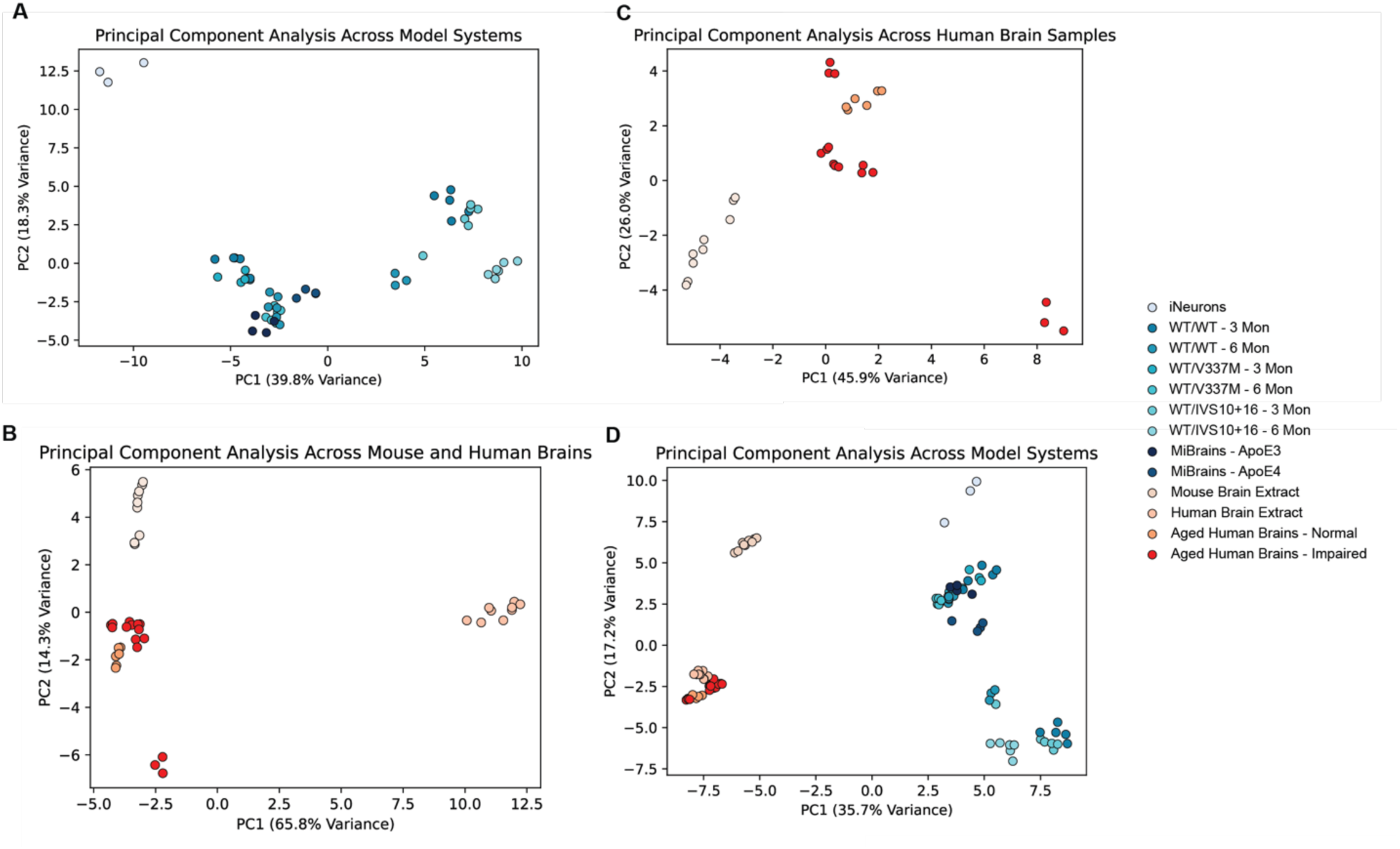
PCA plot of tau proteoforms across cellular (**A**), mouse and human model systems (**B**), clinical samples (**C**) and all systems (**D**). The input for this analysis included only those proteoforms with a mean abundance greater than 1000 ppm across all datasets. Number of replicates are the same as Figure 4B for the model systems: for iNeurons n=3 technical replicates, for WT/WT-3Mon and 6Mon n = 12 (2 donors x 2 conditions x 3 technical replicates), for WT/V337M 3Mon and 6Mon n = 6 (1 donor x 2 conditions x 3 technical replicates), for WT/IVS10+16 3Mon and 6Mon n = 6 (1 donor x 2 conditions x 3 technical replicates), for MiBrains *APOE3* and *APOE4* n=8 (2 x 4 technical replicates), and for Human Brain Extracts and Mouse Brain Extracts n=9 technical replicates each. For Aged Human Brains – Normal n=6 (2 patient samples x 3 technical replicates). For Aged Human Brains – Impaired n=15 (5 patient samples x 3 technical replicates).

**Suppl. Fig. 6:**
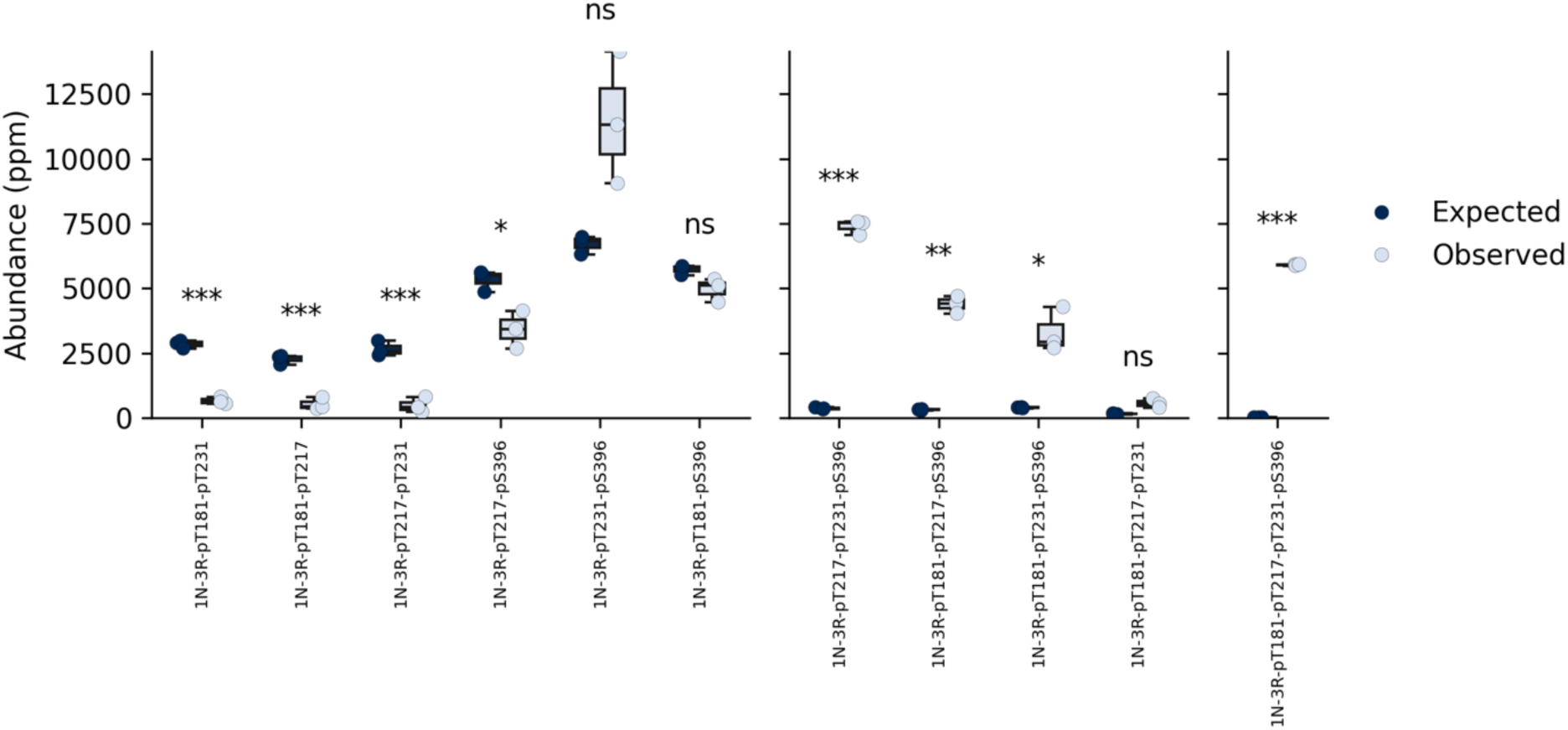
Boxplot of expected vs observed 1N3R proteoform abundances in patient 18. Expected phosphorylation was built with a model assuming that each phosphorylation event occurred independently and was based on the rates of occurrence for each phosphorylation event across all 1N3R molecules (*p<0.05, **p<0.01, ***p<0.001).

